# Gene Flow Creates Fuzzy Species Boundaries in Fence Lizards

**DOI:** 10.64898/2026.04.07.717035

**Authors:** Adam D. Leaché, Hayden R. Davis, Edú B. Guerra, Aracely Herrera, Julio Lemos-Espinal, Matthew K. Fujita, Tanner C. Myers, Sonal Singhal

## Abstract

Species delimitation is a fundamental challenge in systematic biology, particularly for geographically variable taxa with hierarchical population structure and gene flow. Migration-aware coalescent models provide a powerful framework for investigating lineage divergence and accurately defining species boundaries. In this study, we combine statistical evaluations of gene flow with phylogenetic and population structure analyses to delimit species of fence lizards within the *Sceloporus undulatus* complex, a group characterized by extensive population subdivision, mitochondrial DNA introgression, and nuclear gene flow. We find that the *undulatus* complex exhibits uneven variation in genetic, morphological, and bioclimatic traits, resulting in variable distinctiveness among groups. In some cases, species boundaries are recognized by clear genetic discontinuities without gene flow. In others, shallow divergence, paraphyly, and gene flow produce leaky boundaries and fuzzy species limits. Mitochondrial introgression is extensive and concentrated at species boundaries, whereas nuclear gene flow occurs between only a few species and at much lower levels than within species. Neither within-species populations or species are substantially diverged across morphology or bioclimatic space, highlighting the limited utility of these traits for diagnosing species in this group. By integrating estimates of gene flow with phylogenetic and population structure analyses, this study provides a robust and biologically meaningful revised taxonomic framework for the *undulatus* complex that identifies independently evolving lineages as species.

---

Species that are spread across broad geographic areas often consist of multiple populations. Because these populations are often geographically circumscribed, isolated from conspecifics by ecogeographic barriers, and exhibit marked genetic and ecological divergence, they can make attractive candidates for the species level. However, determining the threshold between populations and species is non-trivial. In straightforward cases, variation between species exhibit a level of phenotypic and/or genetic divergence that far surpasses that seen between populations (Hart et al. 2025). In challenging cases, species can exhibit high levels of geographic variation in genetics, morphology, and/or ecology across their range, blurring the distinction between populations and species (Huang 2020; Chambers et al. 2023). A variety of names are used to describe the biological entities that fall into the latter category, including geographically variable species, polytypic species (in cases where a subspecific taxonomy is used), species complexes, superspecies, and taxonomic disaster zones (Prates et al. 2024). Resolving such complex cases is fundamental to systematics and biology more broadly (Barley et al. 2024; Chambers et al. 2025).

One way to unpack such complex cases is by evaluating patterns of gene flow across lineages. Despite their differences, most species concepts, including the biological, evolutionary, and general lineage concepts, coalesce around the idea that species are independently evolving lineages and reproductive communities emerging from the past (Mayr 1942; Mayden 1997; de Queiroz 1998; Maddison and Whitton 2023). Under such concepts, we would expect species to act as independently interbreeding units, and as such, we would expect to see a decrease or complete cessation of gene flow as we cross the population-species boundary (Campillo et al. 2020, Streicher et al. 2024). The evolutionary mechanism thought to drive this association is the evolution of reproductive barriers between species, whether due to local adaptation, sexual selection, and/or genetic drift (Coyne and Orr 2004). Characterizing gene flow, particularly alongside genetic, phenotypic, and ecological divergence, can help clarify the boundaries between populations and species.

Fortunately, new analytical advances allow us to measure gene flow across the divergence continuum and to thus resolve complex evolutionary histories that can only be partially explained using strictly bifurcating models of lineage divergence (Edwards et al. 2016). The multispecies coalescent (MSC) model provides a robust framework for investigating complex demographic histories, including the estimation of effective migration rates among populations and species (Jiao et al. 2021). By allowing some branches of a species tree to evolve under an isolation-migration model while others diverge with no gene flow, the MSC with migration (MSC-M) accommodates both structured populations connected by gene flow and reproductive isolation among species (Flouri et al. 2023). Here, we apply the MSC-M model to understand species boundaries in a geographically variable system: the *Sceloporus undulatus* complex.

## Study System

The *Sceloporus undulatus* species group (“fence lizards”) contains 10 species with a collective distribution spanning the United States and north-central Mexico where they occur in diverse habitats including deserts, prairies, forests, and plateaus (Leaché et al. 2016; Fig. S1). These diurnal species exhibit extensive variation in phenotypic and life history traits, which has made them popular study systems for a wide array of comparative biological studies (Rheubert et al. 2017; Robinson et al. 2024; Robbins and Hegdahl 2024). The main source of taxonomic confusion in the group traces to *S. “undulatus”*, which was previously recognized as a polytypic species with as many as 10 subspecies (Bell et al. 2003). Conflicting signals from ecological, morphological, cytogenetic, and DNA sequence data have confounded attempts to accurately delimit species in *S. “undulatus*” (Cole 1972; Smith et al. 1992; Miles et al. 2002; Leaché and Reeder 2002). A previous phylogenetic analysis of mitochondrial DNA (mtDNA) supported the paraphyly of *S. “undulatus”* with respect to *S. cautus* and *S. woodi,* which resulted in the recognition of *S. “undulatus*” as a species complex consisting of six species: *S. cautus, S. consobrinus, S. cowlesi, S. tristichus, S. woodi,* and *S. undulatus* (Fig. S1; Leaché and Reeder 2002; de Queiroz et al. 2017). Splitting paraphyletic groups, such as this one, into multiple species helps produce a taxonomy that is consistent with phylogenetic relationships. However, reliance on a single genetic locus is error-prone due to incomplete lineage sorting, and particularly in the case of mtDNA, introgression across species boundaries (Petit and Excoffier 2009; Toews and Brelsford 2012; Richmond et al. 2025; Larson et al. 2026). In this study, we use extensive mtDNA and genomic datasets (620 and 445 samples, respectively) to reevaluate species boundaries between the species that comprise the *undulatus* complex: *S. cautus, S. consobrinus, S. cowlesi, S. tristichus, S. undulatus,* and *S. woodi*.

Several previous phylogenetic studies of the *undulatus* complex have uncovered evidence for mtDNA introgression and nuclear gene flow across species boundaries. A study comparing nuclear and mtDNA phylogenies revealed mtDNA introgression at the contact zones between four species pairs (*S. cautus/S. cowlesi, S. consobrinus/S. tristichus, S. consobrinus/S. undulatus, S. cowlesi/S. tristichus*; Leaché 2009); however, sampling was too sparse to estimate population structure or pinpoint species boundaries. A recent investigation of the species boundary between *S. tristichus* and *S. cowlesi* revealed extensive conflict between mtDNA and nuclear data and extensive nuclear paraphyly (Leaché et al. 2025). The discordance between mtDNA and nuclear data in *Sceloporus* seems to stem primarily from gene flow across lineages; thus, accounting for gene flow will both allow us to better reconstruct the evolutionary history of this group and will enable us to define a robust and biologically meaningful taxonomy.

To understand species boundaries in the *Sceloporus undulatus* complex, we conduct phylogeographic analyses across hierarchical levels from populations to species using phylogenetic inference, population structure estimation, and coalescent analyses of migration. Phylogenetic inference is used to identify monophyletic groups as units of analysis, and population structure estimation is used to describe intraspecific variation and detect admixture between groups. Gene flow is then estimated across hierarchical levels using coalescent-based models applied to species tree frameworks. These results are used to identify evolutionary lineages that have a history of durability and a high probability of persisting through evolutionary time. We expect gene flow to be common at the population level, while little to no gene flow is expected between species. We evaluate geographic variation in morphology and ecological niches to provide an integrative perspective on the delimited species and to test whether divergence in these traits corresponds with genetic divergence across the population-species boundary. The outcome is an updated taxonomy for the *undulatus* complex based on extensive geographic sampling and multiple types of data.

## Materials and Methods

### Fieldwork and Sampling

New specimens and samples were obtained for morphological and genetic analysis through both fieldwork and loans from Natural History Museum collections. Fieldwork efforts aimed to sample portions of species ranges that were previously unstudied and at species boundaries. New collections focused on the following regions (and species boundaries): Arizona and New Mexico (*S. cowlesi/S.tristichus*), Colorado (*S. consobrinus/S. tristichus*), Arkansas and Louisiana (*S. consobrinus/S. undulatus*), Northern Mexico (*S. cowlesi/S. consobrinus/S. tristichus*), and Central Mexico (*S. cowlesi/S. cautus/S. exsul*). Tissue loans from Natural History Museums were critical for filling gaps in these and other regions, including Kansas, Kentucky, Nebraska, Oklahoma, Texas, and Wyoming. Detailed voucher specimen information for the mtDNA, nuclear, and morphological datasets is provided in the online Supplementary Information (Tables S1–S3).

### Mitochondrial DNA Genealogy & Delimitation

We collected mtDNA sequence data to increase the geographic resolution of species boundaries and locate regions of potential introgression between species. Genomic DNA was extracted from tissue samples using salt extractions (Aljanabi and Martinez 1997). The mitochondrial *ND1* protein-coding gene (969 bp) was PCR amplified and sequenced using standard methods (Leaché and Cole 2007). New *ND1* sequences were manually aligned (the gene contains no indels) with previously published data to generate a more comprehensive genealogy that included dense geographic and taxonomic sampling (Fig. S1; Table S1).

We estimated a maximum likelihood (ML) phylogeny using IQTREE v.1.6.12 (Nguyen et al. 2015) with 1,000 ultrafast bootstraps (Hoang et al. 2018). The ModelFinder option was used to determine the best-fit nucleotide substitution model on an unpartitioned alignment using the Bayesian information criterion (BIC; Kalyaanamoorthy et al. 2017). Five outgroup species were included, and the most distant outgroup was used to root the tree (*S. graciosus;* Leaché 2010; Wiens et al. 2010; Table S1).

To quantitatively assess mtDNA diversity and focus our mapping of phylogeographic groups, we applied the local minima (“locMin”) method in the R package Spider (Brown et al., 2012). This approach identifies thresholds in the pairwise genetic distance distribution, allowing us to identify meaningful clusters within each species. Analyses were performed separately for the four primary mtDNA species in the *undulatus* complex: *S. consobrinus*, *S. cowlesi*, *S. tristichus*, and *S. undulatus*. To determine the barcode gap threshold, we used the localMinima() function to identify local minima in the pairwise distance distribution, selecting the first minimum occurring after 2% divergence to reduce the influence of shallow intraspecific variation. This method was used to quantify mtDNA diversity and guide the mapping of groups rather than to delimit species boundaries.

### Nuclear DNA Sequencing

We used double-digest restriction site associated DNA sequencing (ddRADseq; Peterson et al. 2012) to obtain a dataset for phylogeographic analyses of both populations and species. The short-read nuclear loci sequenced with this approach can be used directly in multispecies coalescent analyses of gene flow and single nucleotide polymorphisms (SNPs) can be extracted from each locus for population structure inference. We digested 500 nanograms of genomic DNA for 8 hours at 37 °C with 20 units of SbfI (restriction site 5’-CCTGCAGG-3’) and MspI (restriction site 5’-CCGG-3’) in a single reaction with the recommended buffer (New England Biolabs). The fragmented DNA was purified with SeraPure SpeedBeads before ligating barcodes, Illumina adapters, and unique molecular identifiers (UMIs). Pooled samples were selected for fragment sizes between 415 to 515 bp (after accounting for adapter length) on a Blue Pippin Prep size fractionator (Sage Science). The final library amplification used Illumina index primers and high-fidelity Taq polymerase (New England Biolabs). Fragment size distributions and pool concentrations were determined on an Agilent 2200 TapeStation, and qPCR was performed on each library before combining pools for sequencing. A total of 76 pools containing a maximum of 8 samples each were sequenced on four Illumina HiSeq 4000 lanes (50-bp, single-end reads; 43 pools) and one Illumina NovaSeq 6000 lane (100-bp, single-end reads; 22 pools) at the QB3 facility at UC Berkeley and by Novogene Inc. on an Illumina HiSeq X lane (150 bp, paired-end reads; 11 pools). Raw sequencing data from additional samples available on the NCBI Short Read Archive (SRA) were downloaded and combined with the new data (n = 48, Table S2). Approximately 7% of the total sequencing effort was allocated to technical replicates for sample identity verification, with the expectation that replicates would produce identical sequences.

### Nuclear Data Assembly

We assembled the ddRADseq data using iPyrad v.0.7.3 (Eaton and Overcast 2020) with the aid of a reference genome of *Sceloporus tristichus* (Bedoya and Leaché 2021). We de-multiplexed samples using their unique barcode and adapter sequences, allowing no barcode mismatches, and removed UMIs prior to downstream analyses. Sites with phred quality scores < 33 were converted to ‘N’ characters and reads containing five or more low-quality bases were discarded. Reads were clustered using a sequence similarity threshold of 85%, and a minimum base calling depth = 6. Clusters were removed that had an excess of undetermined or heterozygous sites (≥ 5) or too many haplotypes (> 2 for diploids). We processed all samples together and retained the loci that were present for ≥ 50% of the samples. Additional filtering required for downstream analyses are described below.

### Nuclear Data – Broad-scale Analysis

We analyzed the nuclear data using several approaches to identify the major groups to consider for downstream analyses. First, we delineated the major clades supported by the data by estimating phylogenetic relationships with the concatenated nuclear loci (445 samples, 121,189 bp) using IQTREE. The best-fit nucleotide substitution model was selected using the ModelFinder option, and support was measured using 1,000 bootstrap replicates. We rooted the phylogeny with the *cautus-exsul-olivaceus* clade following previous phylogenomic results (Leaché et al. 2016). Second, we examined patterns of genetic variation with principal component analysis (PCA) using the R package dartR v.2.9.7 (Gruber et al. 2018; Mijangos et al. 2022). The PCA results were used to determine which (if any) of the major clades formed discrete clusters. The PCA analysis used a filtered dataset containing one random SNP per locus and a minor-allele frequency threshold (MAF = 0.01). Variant filtering was conducted using VCFtools v0.1.15 (Danecek et al. 2011). Third, we used a network method to infer genealogical relationships, which could include reticulation events that are not modeled explicitly by the concatenated phylogeny. A phylogenetic network was constructed with the NeighborNet algorithm (Bryant & Moulton 2004) in SplitsTree v4.6 (Huson & Bryant 2006) using uncorrected “p” distances calculated from the full alignment (121,189 bp). We mapped clade assignments from IQTREE onto the tips of the network using ‘Network_PieCharts.r’ (Dufresnes et al. 2025) to aid in the visualization of major groups.

### Nuclear Data – Fine-scale Analysis

For each of the major groups identified within the *undulatus* complex, we evaluated fine-scale phylogeographic patterns using population structure inference and species tree analysis. First, to determine the optimal number of populations (*K*) and sample admixture proportions (*Q*), we used the program ADMIXTURE v1.3.0 (Alexander et al. 2009). Population structure was estimated for *K* values ranging from 1–10, and we repeated analyses 10 times with random starting seeds to measure uncertainty in CV scores. The *K* value that provided a combination of a low cross-validation (CV) error and low variation in CV across the 10 replicate runs was selected as the optimal model. We also conducted an analysis with all *undulatus* complex samples combined to estimate population structure and admixture using *K* values ranging from 1–24. We filtered the nuclear loci using VCFtools to select one random SNP per locus and applied a minor allele frequency (MAF) threshold = 0.01. Second, to infer population relationships, we constructed phylogenetic networks using SplitsTree with the concatenated ddRADseq data, and the ADMIXTURE *Q* values were mapped to the tips of the network to visualize groupings. Third, to evaluate whether genetic structure among populations is better characterized as discrete populations or spatial gradients of genetic variation, we compared isolation-by-distance (IBD) patterns within and between populations. Using all variable sites, we estimated F_ST_ for all pairwise individual comparisons within a clade (Reich et al. 2009). We then plotted inverse F_ST_ (F_ST_ / (1 – F_ST_)) against log geographic distance estimated using the R package sf (Pebesma 2018), highlighting any comparisons involving admixed individuals (any individual with >20% probability of ancestry in two or more groups). When there is gene flow between populations, we might expect continuous patterns of IBD within and between populations. However, if populations are experiencing reduced gene flow (i.e., they have evolved reproductive barriers), then IBD will be discontinuous (Good and Wake 1992; Prates et al. 2024).

Finally, we estimated species trees to determine population relationships using the MSC model in BPP v.4.8. (Flouri et al. 2018). We conducted all BPP analyses using the complete ddRADseq loci, incorporating both variant and invariant sites, along with a subset of 1,000 loci without missing data. The optimal population structure model was used to select the number of populations and the sample assignments. The populations were downsampled to include up to eight individuals sampled range-wide to ensure efficient mixing and convergence of the Bayesian analyses. To minimize biases in species tree inference that can be caused by admixed samples, we only retained samples with *Q* values ≥ 0.80. We used inverse gamma priors for population sizes (*θ*) and root age (*τ*_root_) with prior means close to empirical estimates: *θ* ∼ IG(3, 0.002) with mean 0.001 and *τ*_root_ ∼ IG(3, 0.008) with mean 0.004. These priors translate to *θ* ∼ 0.1% sequence divergence within populations and *τ*_root_ ∼ 0.4% sequence divergence for the depth of the species tree. We used a burn-in of 20,000 iterations, and took 400,000 samples, sampling every 2 iterations. Each analysis was repeated four times with different starting seeds and starting trees to assess convergence through the comparison of tree topologies and posterior probability support values.

### Gene Flow Analysis

We used the MSC-M in BPP to estimate migration rates between the major groups supported by the broad-scale analysis and between the populations supported by the fine-scale analyses. We estimated migration rates for all populations with adjacent geographic ranges. We used a stepwise procedure to estimate migration rates and test their significance. The procedure begins with a separate analysis for each bidirectional gene flow event, and then each gene flow even is tested for significance (Ji et al. 2023). The migration rate *M = mN* is measured in the expected number of migrants from the donor population to the recipient population per generation; *N* is the (effective) population size of the recipient population, and m is the proportion of immigrants in the recipient population (from the donor population) every generation, with time moving forward (Flouri et al., 2023). The migration rates were assigned a gamma prior *M ∼* G(1, 10), which assumes an average of 1 migrant individual every 10 generations. The remaining priors and run settings match those used for species tree inference described above. Convergence was assessed by calculating the potential scale reduction factor (PSRF) using the R package CODA (Plummer et al. 2006) with values close to 1 indicating convergence. Posterior distributions from the four separate runs were combined to calculate parameter values and the effective sample size (ESS).

The statistical significance of the estimated migration rates was evaluated using Bayes factors (BF) calculated with the Savage-Dickey density ratio (Ji et al. 2023). The test compares BF_10_ in support of the gene flow model (H_1_) over the null model of no gene flow (H_0_). The Savage-Dickey density ratio requires a cut-off value (ε), which we set to ε = 0.01. If all *M* values in the posterior distribution exceed ε = 0.01, then Bayes factor B_10_ = ∞. The test was considered significant if BF_10_ > 20.

### Morphological Data Analysis

To determine if the widespread species in the *undulatus* complex are supported by morphology, we collected morphometric data and performed multivariate statistical analyses. The sampling locations of specimens included in the morphological analysis (Table S3) were used to assigned them to the main lineages and populations identified in the nuclear phylogeny (Fig. 1). We collected data from nine traits widely used by lizard taxonomists to diagnose species, including five linear measurements and four meristic counts. The linear measurements were snout vent length (SVL) from the tip of the snout to the end of the cloaca; head length (HL) from the tip of the snout to the posterior margin the parietal scale; head width (HW) at the widest point between the sides of the head; femur length (FL) from the articulation with the body to the distal end of the knee joint; and toe length (TL) of the 4^th^ toe from the tip of the claw to where the digit meets the base of the foot. Meristic characters included number of dorsal scales (DS), number of lamellae on the fourth toe (LAM), number of femoral pores (FEMS), and the number of scales in between the femoral pores (FEMSCALES).

**FIGURE 1.**
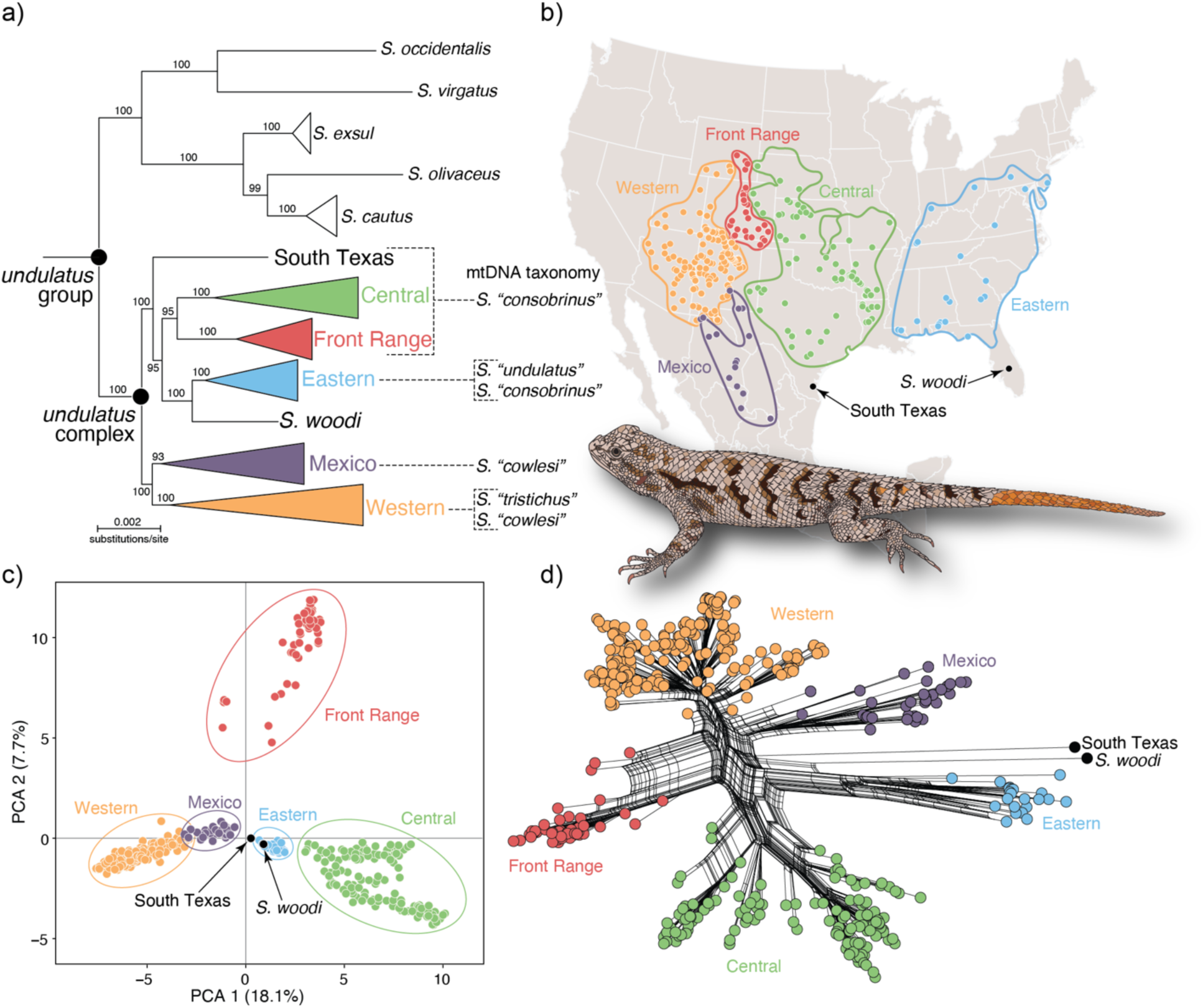
Broad-scale phylogeographic patterns in the *Sceloporus undulatus* species complex using nuclear data. a) Phylogenetic tree of the *undulatus* group (10 species) estimated with IQTREE using 121,189 bp of concatenated nuclear data and 445 samples. All wide-ranging species defined by mtDNA belonging to the *undulatus* complex are paraphyletic. Bootstrap support values are shown on branches. b) Geographic distributions of the major lineages. Sample locations are indicated with dots. c) PCA plot of the SNP data shows clear separation between the Central, Eastern, and Front Range groups, and partial overlap between the Western and Mexico groups. d) Phylogenetic network constructed from the concatenated nuclear data. Artwork of *S. undulatus* by E. Guerra.

Morphometric data analysis was conducted using R v.4.4.2 (R Core Team 2024). First, we analyzed the raw data to assess patterns of variation and sexual dimorphism. Summary statistics were calculated for all traits, and sexual dimorphism was tested using the non-parametric Mann-Whitney U test. Linear models were used to correlate morphological traits with body size (SVL). Second, we performed a principal component analysis (PCA) on the normalized meristic traits using min-max scaling, and on log-transformed continuous traits corrected for body size by extracting residuals from regressions against log-transformed SVL. We assessed the ability of the morphological data to discriminate groups using linear discriminant analysis (LDA) using all measured traits. This method was used to evaluate the ability of the data to correctly classify samples assuming their nuclear DNA-based species assignments. Finally, to test whether morphological divergence scales with genetic differentiation, we calculated population centroids in PCA space and computed pairwise Euclidean and Mahalanobis distances using the stats and MASS R packages. These morphological distances were then compared with average pairwise F_ST_ values.

### Bioclimate Niche Models & Niche Differentiation

We compiled georeferenced occurrence records for the *Sceloporus undulatus* complex (*S. undulatus, S. consobrinus, S. cowlesi, S. tristichus,* and *S. edbelli*) from GBIF, including the synonym “*belli*” to account for widespread use prior taxonomic revisions (Smith et al. 2002). Records lacking coordinates or species-level identification were excluded. Occurrences were assigned to species and populations using minimum convex polygons (MCPs) derived from geographically discrete, monophyletic lineages identified in the nuclear phylogeny. Only records collected after 1970 were retained. To reduce spatial autocorrelation, records were thinned to a minimum distance of 10 km using spThin (Aiello-Lammens et al., 2015; R Core Team, 2025). Calibration areas were defined as MCPs of genetically sampled individuals, with model projections extending to MCP + 200 km. We used 19 bioclimatic variables from WorldClim v2.1 (2.5 arc-min resolution; Hijmans et al., 2005), removing highly correlated predictors (Pearson r > 0.75) and retaining variables with VIF < 10 (Montgomery et al., 2021). To characterize environmental space, we performed a principal component analysis (PCA) on the selected bioclimatic variables using values extracted from 1,000 randomly sampled background points across each calibration area. Occurrence records were then projected onto the resulting PCA axes to visualize differences in environmental space among populations.

Ecological niche models were generated using MaxEnt implemented in ENMTools (Phillips et al. 2006; Warren et al. 2021) with presence data and 1,000 background points sampled within each MCP. We assessed ecological differentiation between populations using pairwise niche identity tests and asymmetric niche similarity tests (Warren et al. 2021). Niche identity tests evaluate whether two ENMs are statistically indistinguishable, indicating that they occupy equivalent environmental space. In contrast, similarity tests compare the observed niche overlap to a null distribution derived from background environments, assessing whether the environments occupied by populations are more similar than expected by chance. These tests were conducted bidirectionally to account for differences in accessible environments. We quantified niche overlap using two metrics: Schoener’s D (Schoener 1968) and Hellinger’s I (Warren et al. 2008). Both values range from 0 (no overlap) to 1 (complete overlap). Finally, we assessed whether ecological nice divergence scales with genetic differentiation by plotting 1 – D (where higher values reflect greater ecological dissimilarity) against average pairwise F_ST_ values.

## Results

### Mitochondrial DNA Genealogy & Delimitation

The final alignment of the mtDNA data contained 620 sequences, and the best-fit nucleotide substitution model selected using the BIC was the TIM+F+I+G4 model. The phylogeny provides strong support (bootstrap ≥ 95%) for the placement of *Sceloporus exsul* and *S. olivaceus* outside of the *undulatus* group (Fig. S1), which contradicts nuclear phylogenomic data (Leaché et al. 2016). Second, the monophyly of *S.* “*cowlesi*” is disrupted by the inclusion of *S. cautus* (Fig. S1), which matches previous studies (Leaché and Reeder 2002).

Species delimitation using the locMin approach identified clear barcoding gaps and multiple phylogeographic groups within each species (Fig. S2): *S. undulatus, n* = 3 (threshold = 4.02%), *S. consobrinus, n* = 6 (threshold = 3.12%), *S. cowlesi, n* = 13 (threshold = 2.27%), and *S. tristichus, n* = 13 (threshold = 2.62%). Detailed phylogeographic results of each species with species delimitation results mapped to the phylogeny are provided in Figures S3–S6.

### Nuclear Data – Broad-scale Analysis

The ddRADseq dataset produced 3,896 loci for 445 samples (Table S4), and the concatenated dataset contains 121,189 base pairs. The best-fit nucleotide substitution model selected using the BIC is the GTR+F+R4 model. The nuclear phylogeny places *S. cautus* outside of the *undulatus* complex (Fig. 1a), which contradicts the mtDNA genealogy, but is concordant with previous phylogenetic analyses of nuclear data (Leaché et al., 2016). The phylogeny splits the *undulatus* complex into seven groups, and all wide-ranging species defined by mtDNA are paraphyletic (Fig 1a; *S.* “*consobrinus*”*, S.* “*cowlesi*”*, S.* “*tristichus*”, and *S.* “*undulatus*”). The nuclear phylogeny splits *S.* “*consobrinus*” into three clades: Central = populations from throughout the central U.S, Front Range = populations from the Front Range of Colorado and central Wyoming, and South Texas = a population from Kenedy Co., Texas (Fig. 1a,b). A broadly distributed Western clade contains populations of *S. “cowlesi”* and *S. “tristichus*” from throughout the Desert Southwest and Mexico (Fig. 1a,b). Population structure analysis of the *undulatus* complex using PCA (436 samples, 3,718 SNPs) supports clusters that correspond to the major groups supported by the phylogeny (Fig. 1c). The PCA plot separates the Central, Eastern, and Front Range groups, but the Western and Mexico groups overlap partially (Fig. 1c). The phylogenetic network (Fig. 1d) also groups samples in accordance with the phylogeny and the PCA plots.

### Nuclear Data – Fine-scale Analysis

Population structure analyses using ADMIXTURE support multiple populations within the major clades of the *undulatus* complex (Fig. 2; Fig. S7). The best-fit model for the Eastern clade is *K* = 2 (Fig. 2b, Fig. S8). The populations are split between east and west with no evidence for admixed samples at the population boundary (Fig. 2b, Fig. S8). However, one divergent sample from Georgia positioned at the base of the phylogeny is admixed (Fig. S8). A model with *K* = 3 was selected for the Mexico clade (Fig. 2c); this model had low variation among replicate runs despite the *K* = 2 model having a lower average CV score, but high variation (Fig. S7). The three populations include northern and southern Mexico groups that are admixed where they meet in Chihuahua, and a third population with samples in Texas and New Mexico (Fig. 2; Fig. S9). The best-fit model for the Front Range clade is *K* = 3 with populations in the northern, southeast, and southwest portions of the range (Fig. 2d). Admixed samples are found at the geographic boundaries separating the populations (Fig. 2d, Fig. S10). The best-fit model for the Central clade is *K* = 5 (Fig. 2e, Fig. S11). The five populations form a ring-like distributional pattern with admixed samples found at each of the connections between adjacent populations (Fig. 2e, Fig. S11). The best-fit model for the Western clade is *K* = 6 (Fig. 2f, Fig. S12). All populations show extensive admixture (Fig. 2f, Fig. S12). Phylogenetic networks cluster samples similarly to the phylogeny, and place admixed samples in intermediate positions (Figs. S8-12).

**FIGURE 2.**
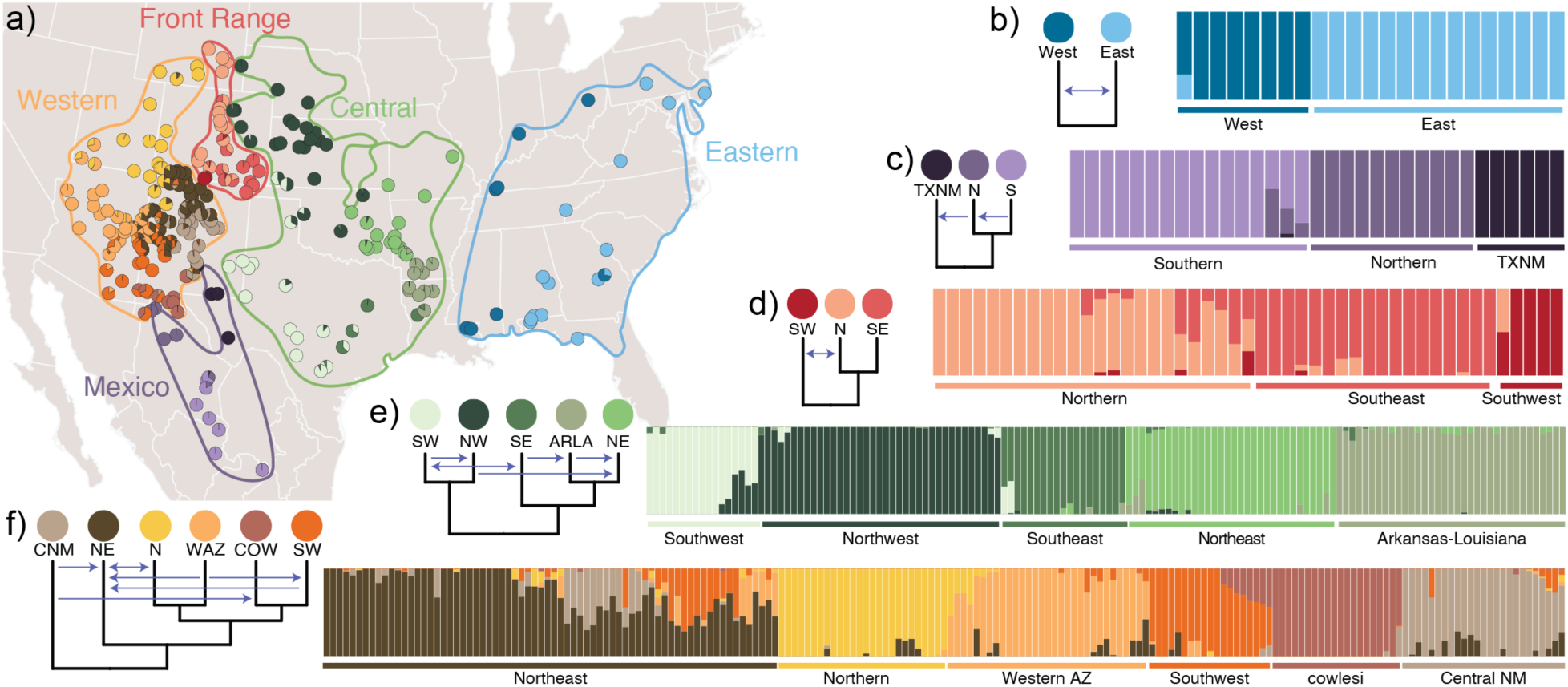
Phylogeography of the widespread lineages in the *undulatus* complex using ddRADseq data. a) Geographic distributions with pie-charts showing sample admixture proportions. Samples ≤ 10km apart are merged. b–f) Bar plots showing population structure, and species trees with arrows indicating significant migration estimates.

Estimating population structure using all *undulatus* complex samples combined produces ambiguous results for the optimal *K* value, which is nearly equivalent for *K* values ranging from 17–24 (Fig. S7). Assuming *K* = 19, which equals the total number of populations from the separate analyses described above, supports admixture between the Western clade with both Mexico and the Front Range (Figs. S13–S14). Isolation by distance (IBD) is recovered across all comparisons (P-values ≤ 0.05; Table S5), with genetic similarity generally declining as geographic distance increases, although the strength of this relationship varies among populations (Figures S15–S18). Within-population IBD patterns are generally weaker compared to between-population comparisons, which exhibit steeper differentiation that reflect population structure at broader spatial scales.

### Gene Flow Analysis

Migration rates estimated using the MSC-M model in BPP are provided in Table S6. Convergence diagnostics indicate that the replicate analyses reached stationarity with PSRF scores close to 1.0 and combined ESS values ≥ 200 (Table S7). Three gene flow events are significant (BF_10_ > 20) in the analyses of gene flow among the major *undulatus* complex clades, including migration from Mexico into the Western clade, and migration into the Front Range from the Central and Western clades (Fig. 3). All other migration events between species with adjacent geographic distributions are rejected (BF_10_ < 20; Table S6). The highest migration rate estimated at the broad-scale is from Mexico → Western with a mean *M* = 0.1678 (95% HPD = 0.0437–0.2924; Table S6). This migration rate is exceeded within each clade by at least one population-level migration rate (Fig. 3). Bidirectional migration is supported within the Eastern clade (Fig. 3). In the Mexico clade, two unidirectional migration rates are significant and suggest a pattern of northward migration (Fig. 3). Within the Front Range, bidirectional migration is supported between northern and southwest populations with the higher rate from northern → southwest with a mean *M* = 1.0276 (95% HPD = 0.7044–1.3643; Fig. 3; Table S6). Six migration rates are significant in the Central clade with the highest rate from southwest → southeast with a mean *M* = 0.530 (95% HPD = 0.2645–0.8034; Fig. 3; Table S6). Seven migration rates are significant in the Western clade, and the mean migration rates range from 0.1152–0.6487 (Fig. 3; Table S6).

**FIGURE 3.**
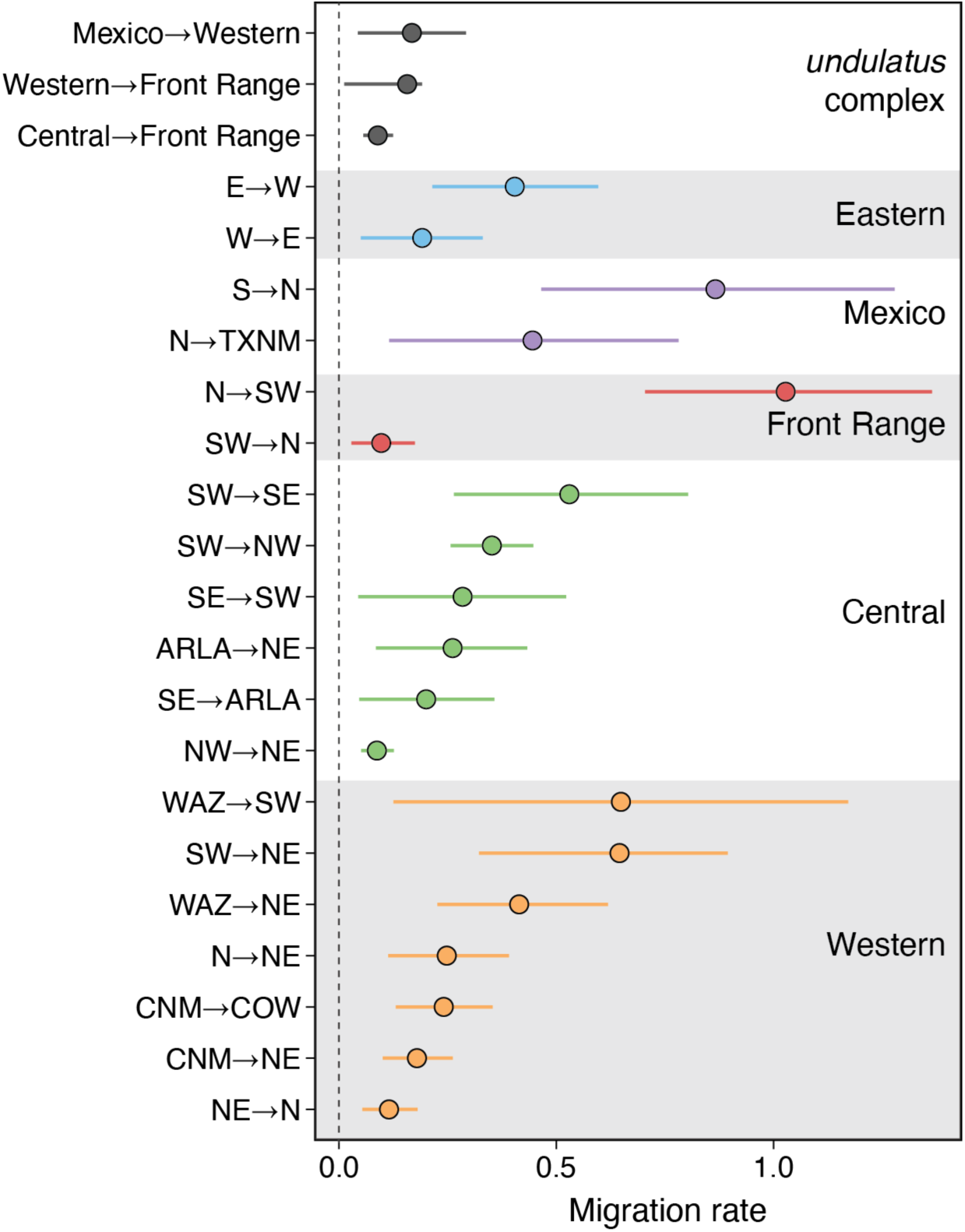
Migration rates in the *Sceloporus undulatus* complex estimated with the nuclear data using the MSC-M model in BPP. The top three comparisons are between major *undulatus* complex clades; all other comparisons are between populations within each clade. Only significant migration rates (BF_10_ > 20) are shown.

There is a clear negative relationship between migration rate and genetic differentiation, with higher F_ST_ values associated with lower migration rates (Fig. 4a). Population comparisons show a wider range of migration rates at lower to moderate F_ST_ values, while broad-scale comparisons cluster at higher F_ST_ values with consistently low migration rates (Fig. 4a). Within the broad-scale comparisons, the Western–Mexico pair stands out with a relatively high migration rate and low F_ST_ value, positioning it at the upper end of population comparisons (Fig. 4a). Overall, these patterns indicates that greater genetic divergence corresponds to reduced gene flow.

**FIGURE 4.**
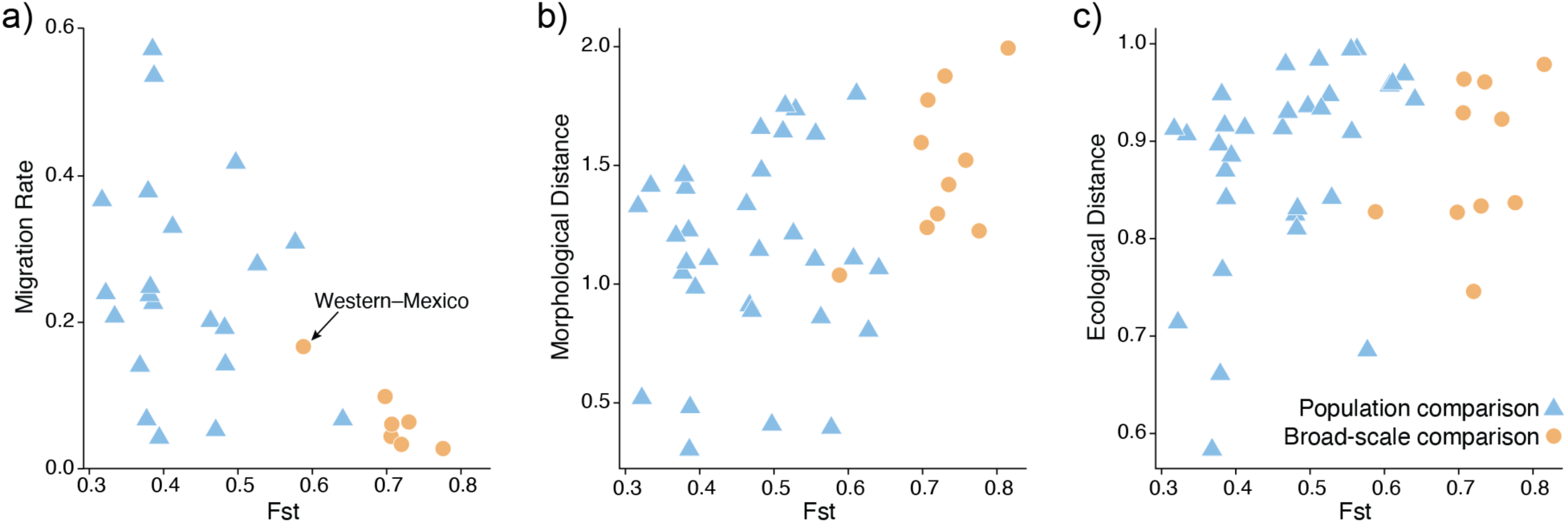
Patterns of morphological, ecological, and genetic divergence in the *Sceloporus undulatus* complex. Genetic differentiation as measured by F_ST_ is compared against a) migration rate, b) morphological distance, and c) ecological distance. All comparisons are colored by the comparison type (population comparisons = blue triangles; broad-scale clade comparisons = orange circles).

### Morphological Data Analysis

The raw morphological data and a summary of trait variation are provided in Tables S8–S9. Mann-Whitney U tests provided idiosyncratic results for which traits showed significant sexual dimorphism, indicating that sexual dimorphism may not be relevant for these traits (Table S10). The Western clade exhibits significant sexual size dimorphism (SVL; *p*-value = 8.8E-07; Table S10). Linear models showed strong correlations between most traits and SVL (Fig. S19), with the exceptions of LAM (*p* = 0.163) and FEMSCALES (*p* = 0.929). PCA of the morphological data does not provide clear clustering of genetic groups (Fig. S20). The first three components account for 60% of the variation (Table S11). The LDA analysis provides the highest classification success for the Front Range samples (92.9%), and all misclassifications belong to the Western clade (Table S12). The Western clade has the lowest classification success (48.1%; Table S12).

We find that morphological distance tends to increase with F_ST_, indicating that genetic differentiation scales with morphological divergence (Fig. 4b). The population comparisons cover a wider range of lower to moderate F_ST_ values and morphological distances, while the broad-scale comparisons are clustered at higher F_ST_ values and generally higher morphological distances (Fig. 4b). This pattern suggests that morphological divergence aligns broadly with genetic divergence, with stronger differentiation observed at broader scales.

### Bioclimate Niche Models & Niche Differentiation

Maxent models performed well for the *S. undulatus* complex. AUC values ranged from 0.696 to 0.983 (mean = 0.854 ± 0.077 SD; Table S12). Across models, the most important bioclimatic predictors were temperature seasonality (Bio4), mean temperature of the driest quarter (Bio9), precipitation seasonality (Bio15) (Table S12). Ecological niche models for all focal populations are provided in Figures S21–S26. Overall, model performance was robust and suitable for subsequent niche overlap analyses. Niche identity tests revealed varying degrees of overlap across most comparisons (Table S13). At the broad-scale, niche identity values were consistently low (e.g., Schoener’s D = 0.021–0.254; I = 0.096–0.531), indicating substantial ecological divergence among clades. The highest niche overlap occurred between the Central and Western clades (D = 0.254, I = 0.531), while Front Range and Eastern clades exhibited the least overlap (D = 0.021, I = 0.096). Complementarily, population-level comparisons support a wide range of overlap, from very low values (e.g., Central clade ARLA vs. SW; D = 0.004, I = 0.027) to relatively high values (e.g., Western clade CNM vs. NE; D = 0.416, I = 0.670), suggesting a broad gradient of ecological differentiation among populations.

Asymmetric similarity test results revealed further insights (Table S13). At the broad-scale, several pairs showed significant niche divergence in both directions (e.g., Central vs. Mexico clades), while others were significant in one direction (e.g., Central vs. Western clades significant only in the 1 ← 2 direction). Such asymmetry suggests that one group occupies a broader or more environmentally variable background, while the other has a more constrained niche. In these cases, the broader-niched species may find suitable conditions in the range of the other, but not vice versa. Population comparisons often showed asymmetrical results, where significant niche similarity was detected in one direction but not the other (e.g., Eastern clade east vs. west).

Ecological distances are generally high across most F_ST_ values, and population comparisons exhibit a broader range of ecological distances at lower F_ST_ values (Fig. 4c). Broad-scale comparisons tend to cluster at higher F_ST_ values compared to population comparisons (Fig. 4c). Overall, these patterns suggest that ecological divergence is generally high across regardless of genetic differentiation.

## Discussion

Integrating statistical models for evaluating gene flow with phylogenetic and population structure inferences provides a useful framework for delimiting species with structured populations, gene flow, and mitochondrial introgression. Since phylogenies and population structure capture different levels of evolutionary divergence, applying this framework across these hierarchical scales provides a robust assessment of the populations that warrant recognition as species and the resulting taxonomic implications. Within the *Sceloporus undulatus* complex, this approach suggests a continuum of divergence, which we can see across patterns of gene flow, morphological divergence, and ecological divergence.

Gene flow within the *undulatus* complex is substantially higher between populations within the same species than between different species, highlighting an important distinction along the population–species continuum. Specifically, coalescent-based analyses of gene flow reveal significant migration between populations at relatively high levels, often exceeding one migrant per generation (Fig. 3), indicating substantial genetic connectivity within species. In contrast, the few significant migration events detected between species occur at much lower rates, typically below 0.2 migrants per generation, suggesting stronger reproductive isolation. These patterns provide insight into how gene flow shifts across stages of evolutionary divergence, with migration rates declining as lineages become more genetically and evolutionarily distinct (Fig. 4a). In parallel, niche comparisons indicate substantial environmental divergence in both population and brad-scale comparisons (Fig. 4c). Together, these patterns are consistent with divergence across ecological gradients, where environmental differences may contribute to population differentiation despite ongoing connectivity. This gradient in gene flow has important taxonomic implications, suggesting that the observed reduction in gene flow reflects the emergence of independently evolving lineages across environmentally distinct conditions.

Although morphological divergence is relatively modest across both populations and species in the *undulatus* complex (Fig. 4b), species appear to exhibit greater morphological divergence than populations, and morphological divergence increases with genetic divergence (Fig. 4b). This same pattern is recapitulated for bioclimatic divergence (Fig. 4c). Across all three metrics of gene flow, morphology, and bioclimatic niche, divergence is relatively continuous across populations to species. Although species clearly exhibit greater divergence than populations, there is no obvious gap that allows us to clearly demarcate the population-species threshold.

In some cases, species boundaries align with clear discontinuities in genetic variation, where gene flow is effectively absent among species but persists among populations within species. In other cases, however, structured populations exhibit shallow divergence, paraphyly, and ongoing or historical gene flow, obscuring species limits and creating ambiguity in taxonomic assignment. These more complex scenarios highlight the limitations and risks of relying on any single line of evidence for taxonomic inference.

### Mitochondrial DNA introgression

Mitochondrial DNA introgression localized at species boundaries is common within the *undulatus* complex and is a primary source of taxonomic confusion (Leaché 2009). As a result, recognizing mtDNA clades as species is problematic, because genealogical patterns in mtDNA can reflect introgression rather than true evolutionary independence. In contrast, multilocus nuclear data provide a reliable basis for species delimitation that also offer important insight into patterns of gene flow among populations and species. Evidence of mtDNA introgression in the *S. undulatus* group is found at both deep and shallow phylogenetic levels. At the deepest level, alternative placements of *S. olivaceus* and *S. exsul* either inside the species group (nuclear DNA) or outside of it (mtDNA) point to either incomplete lineage sorting or ancient introgression with undescribed ghost lineages. At shallow levels, species boundaries that seem sharp and precise based on mtDNA data are non-existent with nuclear DNA, as is the case with the “invisible” species boundary that separates the mtDNA clades described as *S.* “*tristichus*” and *S.* “*cowlesi*”.

The natural history of these species offers some clues to why conflicting nuclear and mtDNA phylogenetic patterns might be expected in the *undulatus* complex. Adult males are territorial and known for displaying their blue abdominal and gular regions while performing push-up and head-bob behaviors that serve as social signals in territory disputes, courtship, and species recognition (Martins 1993; Hews et al. 2011). Territorial males adjust their home range size to overlap with multiple females, while females maintain smaller ranges, resulting in a significant sex-based difference in home range size (Ferner 1974; Haenel et al. 2003). Female philopatry can influence the spatial distribution of sex-linked loci and cause biases in species delimitation (Eberle et al. 2019). Therefore, because of male biased dispersal in *Sceloporus*, we expect nuclear DNA to exhibit more gene flow and relatively weaker geographic structure compared to maternally inherited mtDNA (Toews and Brelsford 2012). Additionally, the larger effective population size of nuclear loci should result in slower coalescent times compared to mtDNA and produce less decisive genealogical results (Palumbi et al. 2001). Contrasting mtDNA and nuclear phylogeographic patterns reveals that mtDNA alone has misrepresented species boundaries in the *undulatus* complex, whereas nuclear loci provide a robust framework for species delimitation by more accurately reflecting both gene flow and evolutionary history.

### Taxonomic Recommendations

The current taxonomy for the *Sceloporus undulatus* complex is largely based on mtDNA. This taxonomy is effectively invalid because, as shown by our multilocus nuclear data, it does not reliably capture the evolutionary history of the complex. Here, we propose taxonomic revisions that accurately reflect both the evolutionary history of the group and incorporate characterizations of patterns of gene flow and ecological and bioclimatic divergence (Fig. S27). This taxonomic framework thus establishes a more stable and biologically meaningful taxonomy by aligning taxonomic units with independently evolving lineages (de Queiroz 2007; Maddison and Whitton 2023).

### Western clade

The sharp boundary between *S. cowlesi* and *S. tristichus* observed with mtDNA that extends across central Arizona and New Mexico is absent from the nuclear data, which instead support a boundary in northern Mexico between Western and Mexico clades (Fig. 1; Leaché et al. 2025). The type localities of *S. tristichus* and *S. cowlesi* both belong to the Western clade, and *Sceloporus tristichus* Cope in Yarrow, 1875 was described first and has priority. The type locality of *S. tristichus* is “Taos, Taos Co., New Mexico”. Therefore, under the taxonomic framework of the nuclear phylogeny, *Sceloporus cowlesi* is a synonym of *S. tristichus.* The six populations described within *S. tristichus* are connected by broad zones of admixture (Fig. 2) and exhibit continuous patterns of isolation-by-distance between populations (Fig. S18). These population boundaries are fuzzy, as might be expected for weakly differentiated populations that are at early stages of divergence (Leaché et al. 2025). As such, no within-species populations should be elevated to species level.

### Mexico clade

The sister group to *S. tristichus* is the Mexico clade, which is distributed throughout Mexico and into West Texas and southern New Mexico (Fig. 2). *Sceloporus edbelli* Smith, Chiszar and Lemos-Espinal, 2002 is available and takes priority for populations distributed in Mexico from northwestern Chihuahua to southern Coahuila, northwestern Zacatecas, and northeastern Durango. The type locality is “two mi S León Guzmán, Durango, Mexico”. Populations belonging to *S. edbelli* occur in sympatry with two other species in the *undulatus* group, which further supports the recognition of this species despite the limited gene flow that is detected with *S. tristichus* (Fig. 3). At the northern end of their distribution, *S. edbelli* and *S. tristichus* co-occur at two locations in Chihuahua, Mexico: the Cerro Colorados near the southern end of Samalayuca Sand Dunes and the foothills of Sierra de San Luis (Pradera de Janos). In the south, *S. edbelli* occurs in sympatry with *S. cautus* in Zacatecas near Anáhuac. The boundaries between the populations in *S. edbelli* are sparsely sampled, but the relatively high levels of gene flow between populations suggests that they should not be elevated to species.

### Central clade

*Sceloporus consobrinus* Baird and Girard, 1853 is retained for the Central clade that includes the type locality of Quartz Mountain State Park, Kiowa County, Oklahoma (see Bell and Smith 2000). The species has a broad distribution with at least five populations distributed from eastern Wyoming to the Mississippi River (Fig. 2); none of these populations exhibit substantial enough divergence to be elevated to species level.

### Front Range clade

*Sceloporus erythrocheilus* is proposed for populations distributed throughout the Front Range of Colorado from Wyoming to northeastern New Mexico (Fig. 2). This taxon was originally described as *Sceloporus undulatus erythrocheilus* Maslin, 1956 with a type locality “Nineteen mi E Model, Purgatoire River, Las Animas County, Colorado”. Three populations are supported in *S. erythrocheilus* (Fig. 2), none of which should be considered species given their low divergence and substantial admixture. Low levels of gene flow from *S. tristichus* and *S. consobrinus* are detected in *S. erythrocheilus* (Fig. 3).

### Eastern clade

*Sceloporus undulatus* is retained for the Eastern clade, but with a modified western boundary that is shifted to the west to align with the Mississippi River. The separation between *S. undulatus* and *S. consobrinus* corresponds with the Mississippi River, but more extensive sampling is needed to test the permeability of this boundary. Several populations in the western portion of the range of *S. undulatus* carry mtDNA haplotypes belonging to *S. consobrinus,* yet no significant nuclear gene flow was detected between these species. Increased sampling is needed to refine the boundary between populations within *S. undulatus*.

### South Texas

*Sceloporus thayerii* Baird and Girard, 1852 is tentatively applied to the South Texas lineage (Fig. 2), which aligns geographically with part of the type locality, “Indianola [Calhoun, County, Texas]” (Bell et al. 2003). However, this assignment should be considered provisional, as the species is represented in our study by a single specimen. More extensive sampling in South Texas is needed to determine the geographic extent and taxonomic validity of this putative species.

### Future Directions

The *Sceloporus undulatus* complex exhibits a continuum of genetic divergence across species and populations, produced by uncertain species boundaries with signatures of gene flow and mitochondrial DNA introgression. Divergence in morphology and ecology also spans a broad spectrum with increased differentiation among species compared to populations. These results help identify geographic regions where diversity is shaped by gene flow and mitochondrial DNA introgression. The shallowest evolutionary divergence is located at the boundary between *S. tristichus* and *S. edbelli;* these species exhibit the highest level of gene flow and least genetic differentiation measured between any species pair (Fig. 4a). Mitochondrial introgression is detected between five additional species pairs (Fig. S28): 1) *S. tristichus* haplotypes are found in southwestern populations of *S. erythrocheilus* in southern Colorado, 2) *S. erythrocheilus* haplotypes occur in *S. consobrinus* in eastern Colorado and western Kansas, 3) *S. consobrinus* haplotypes are found on the eastern side of the Mississippi River in western populations of *S. undulatus,* 4) *S. thayerii* has a haplotype belonging to *S. consobrinus* in southern Texas, and 5) introgression between *S. cautus* and *S. edbelli* occurs in both directions in Zacatecas. Targeted studies of these evolutionary hotspots will be key to understanding population-species boundary formation in the *undulatus* complex.

The morphometric dataset provided little resolution for distinguishing among the genetic groups. Trait variation was high within groups with extensive overlap among them, obscuring any consistent diagnostic differences. This pattern was further supported by the PCA results, where different lineages clustered together without clear separation. Overall, despite high levels variation in morphometric and meristic traits, they appear to be poor predictors of the underlying genetic structure in this system, suggesting that phenotypic variation is weakly linked to lineage divergence, possibly due to strong environmental and/or plastic responses. For example, in *S. undulatus*, analyses of full genomes support adaptive responses in limb length to selective pressures imposed by invasive fire ants (Assis et al. 2025), indicating that morphological variation has a genetic basis shaped by local adaptation, rather than directly reflecting patterns of population divergence. Several key diagnostic traits used to distinguish taxa within the *undulatus* complex are adult male coloration, particularly differences in the extent and patterning of abdominal and gular patches. Coloration differences such as these could be more informative for distinguishing lineages since they are associated with reproductive strategies and/or mating preferences in Phrynosomatid lizards (Corl et al. 2026). Future studies would benefit from quantitatively assessing the genetic basis of ecological, morphological, and color pattern traits in the *undulatus* complex and explicitly linking these differences to local adaptation and evolutionary divergence at population and species boundaries.

## Supplementary Material

Data to be made available on the publishers website.

## Acknowledgements

We thank X anonymous reviewers, the Associate Editor (X), and the Editor (X) for their constructive comments on the manuscript. We appreciate the contributions of specimens from Natural History Museums: Lauren Sheinberg, Jens Vindum, and Rayna Bell (California Academy of Sciences), Carol Spencer and Jim McGuire (Museum of Vertebrate Zoology), Greg Pauly and Neftali Camacho (Natural History Museum Los Angeles County), William Stark and Curtis Schmidt (Sternberg Museum of Natural History), Jessa Watters and Cameron Siler (Sam Noble Oklahoma Museum of Natural History), Toby Hibbitts and Lee Fitzgerald (Texas Cooperative Wildlife Collection), Eli Greenbaum, Carl Lieb, and Vicky Zhuang (University of Texas at El Paso Biodiversity Collections), Sharon Birks and Peter Miller (Burke Museum of Natural History and Culture), Wendy Estes-Zumpf and James Cash (Wyoming Game and Fish Dept.), David Laurencio, Jamie Oaks, and Daniel Warner (Auburn University Museum of Natural History), and Joe Ehrenberger (Adaptation Environmental Services). We also thank Jeremy Chamberlain, Donavan Jackson, and André Carvalho for assisting with the collection of specimens We thank state agencies for scientific collecting permits, including Arizona Game and Fish, New Mexico Game and Fish, and Colorado Parks and Wildlife, Colorado State Land Board, Boulder County Parks and Open Spaces.

## Data Availability

Mitochondrial DNA sequences are deposited at NCBI Genbank (XXXX-XXXX). Demultiplexed ddRADseq data are deposited at NCBI SRA (PRJNA XXXX). Data available from the Dryad Digital Repository: https://doi.org/10.5061/dryad.547d7wmnx.

