## Supplemental Figures for "Gene Flow Creates Fuzzy Species Boundaries in Fence Lizards"

**Table of contents**

**FigS1.** Current taxonomy based on mitochondrial DNA.
**FigS2.** Species delimitation using the mtDNA data.
**FigS3.** Phylogeographic patterns within *Sceloporus cowlesi* using mtDNA.
**FigS4.** Phylogeographic patterns within *Sceloporus consobrinus* using mtDNA.
**FigS5.** Phylogeographic patterns within *Sceloporus undulatus* using mtDNA.
**FigS6.** Phylogeographic patterns within *Sceloporus tristichus* using mtDNA.
**FigS7.** Admixture analysis results and cross-validation (CV) error scores using SNP data.
**FigS8.** Phylogeographic patterns within the Eastern clade using nuclear data.
**FigS9.** Phylogeographic patterns within the Mexico clade using nuclear data.
**FigS10.** Phylogeographic patterns within the Front Range clade using nuclear data.
**FigS11.** Phylogeographic patterns within the Central using nuclear data.
**FigS12.** Phylogeographic patterns within the Western clade using nuclear data.
**FigS13.** Admixture results ($K$ = 19) for the broad-scale analysis.
**FigS14.** Admixture results ($K$ = 19) for the broad-scale analysis, zoomed in on the Western USA.
**FigS15.** IBD results for the Eastern and Mexico populations.
**FigS16.** IBD results for the Front Range populations.
**FigS17.** IBD results for the Central populations.
**FigS18.** IBD results for the Western populations.
**FigS19.** Morphology data scatterplots of trait correlations with body size.
**FigS20.** Morphology data PCA plots.
**FigS21.** Ecological niche models for the Eastern clade.
**FigS22.** Ecological niche models for the Mexico clade.
**FigS23.** Ecological niche models for the Front Range clade.
**FigS24.** Ecological niche models for the Central clade.
**FigS25.** Ecological niche models for the Western clade.
**FigS26**. Ecological niche model PCA plots for each species and clade.
**FigS27.** Proposed taxonomic changes for the *Sceloporus undulatus* complex.
**FigS28.** Summary of mitochondrial introgression in the *Sceloporus undulatus* complex.

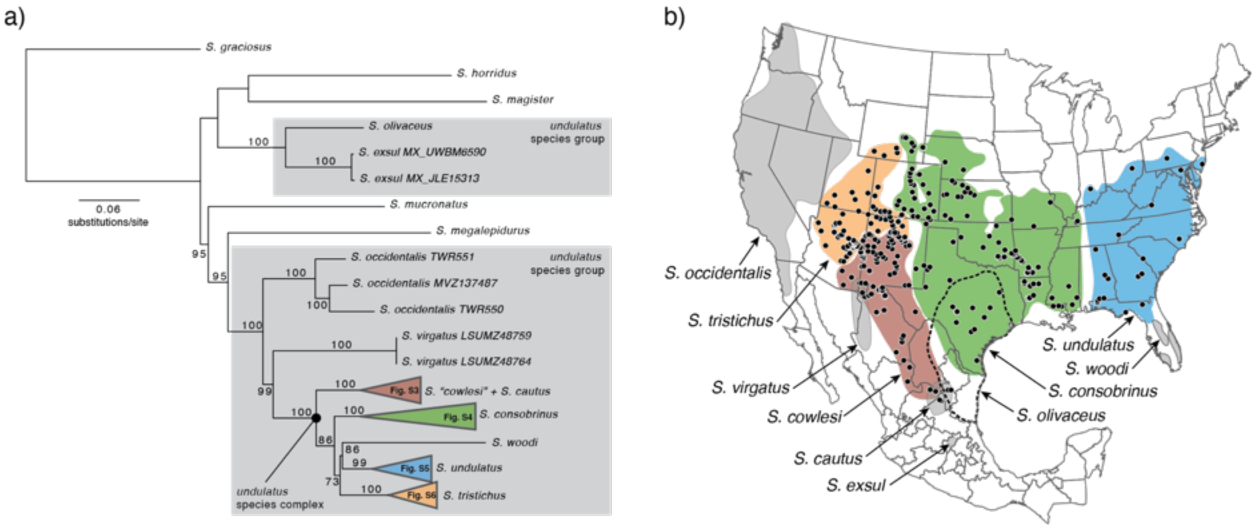


FIGURE S1. Current taxonomy and species boundaries for the *Sceloporus undulatus* species group based on mtDNA sequence data. a) Phylogeny inferred from the *ND1* protein-coding gene using IQTREE. Branches are labeled with bootstrap support values. Clade specific maps are shown in Figures S3–S6, as indicated in the phylogeny. b) Species distributions and sampling of the widespread species in the *undulatus* complex (black dots).

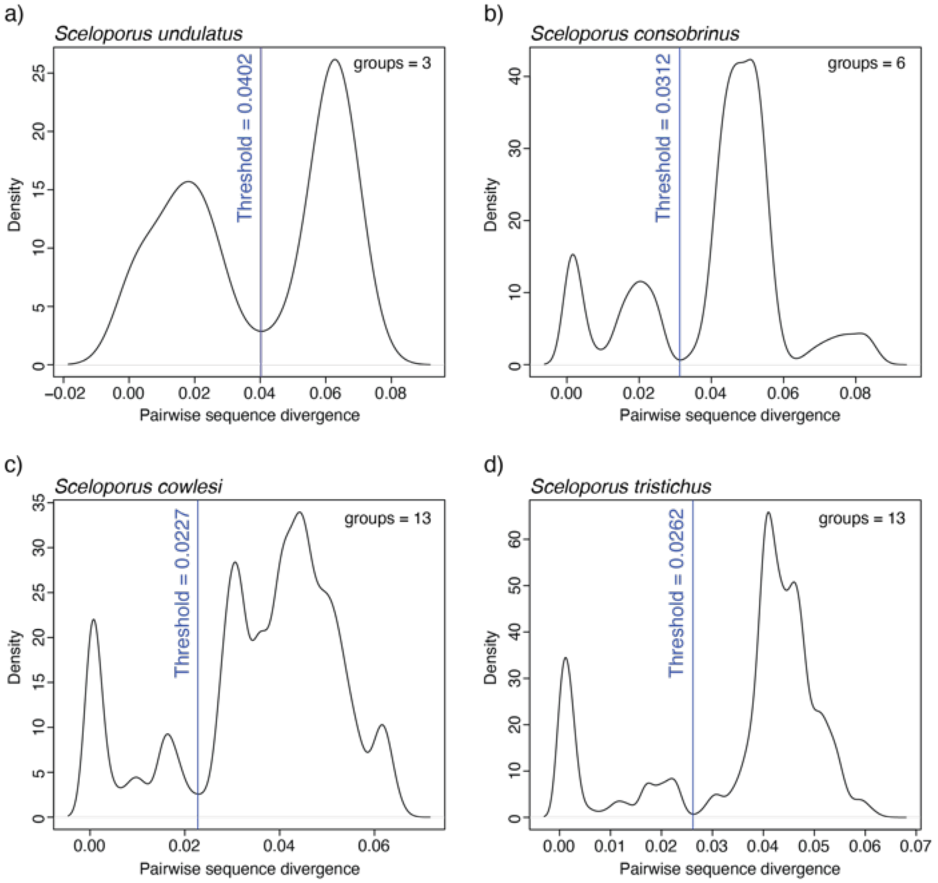


FIGURE S2. Species delimitation using the mtDNA sequence data with the locMin method implemented in the R package Spider for a) *S. undulatus,* b) *S. consobrinus,* c) *S. cowlesi*, and d) *S. tristichus*. The first local minimum found after 2% divergence was selected as the barcoding gap threshold.

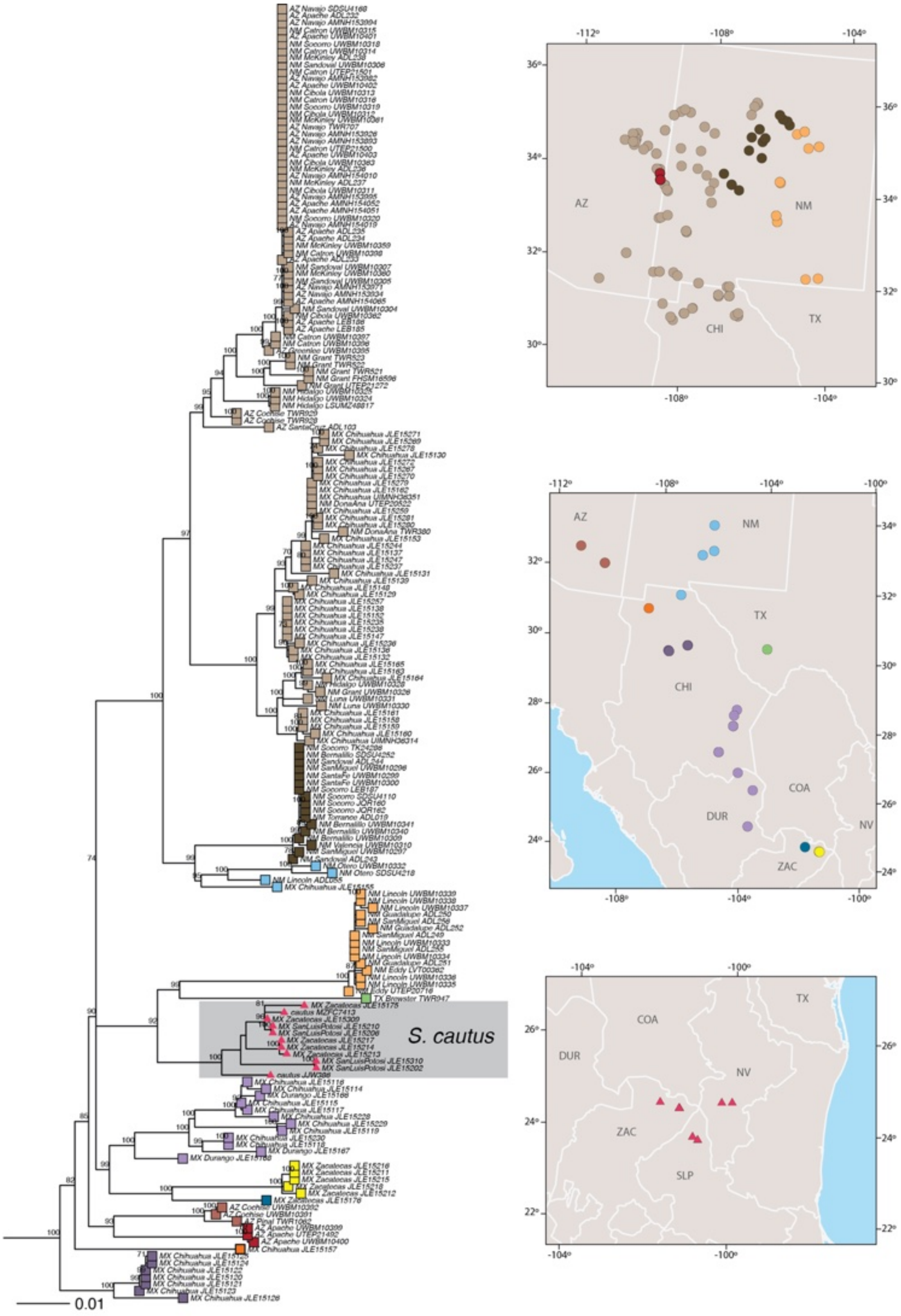


FIGURE S3. Phylogeographic patterns within *Sceloporus cowlesi* and *S. cautus* inferred using mtDNA. The phylogeny was estimated with IQTREE. Tips with the same colors reflect species delimitation results from the locMin method implemented in the R package Spider.

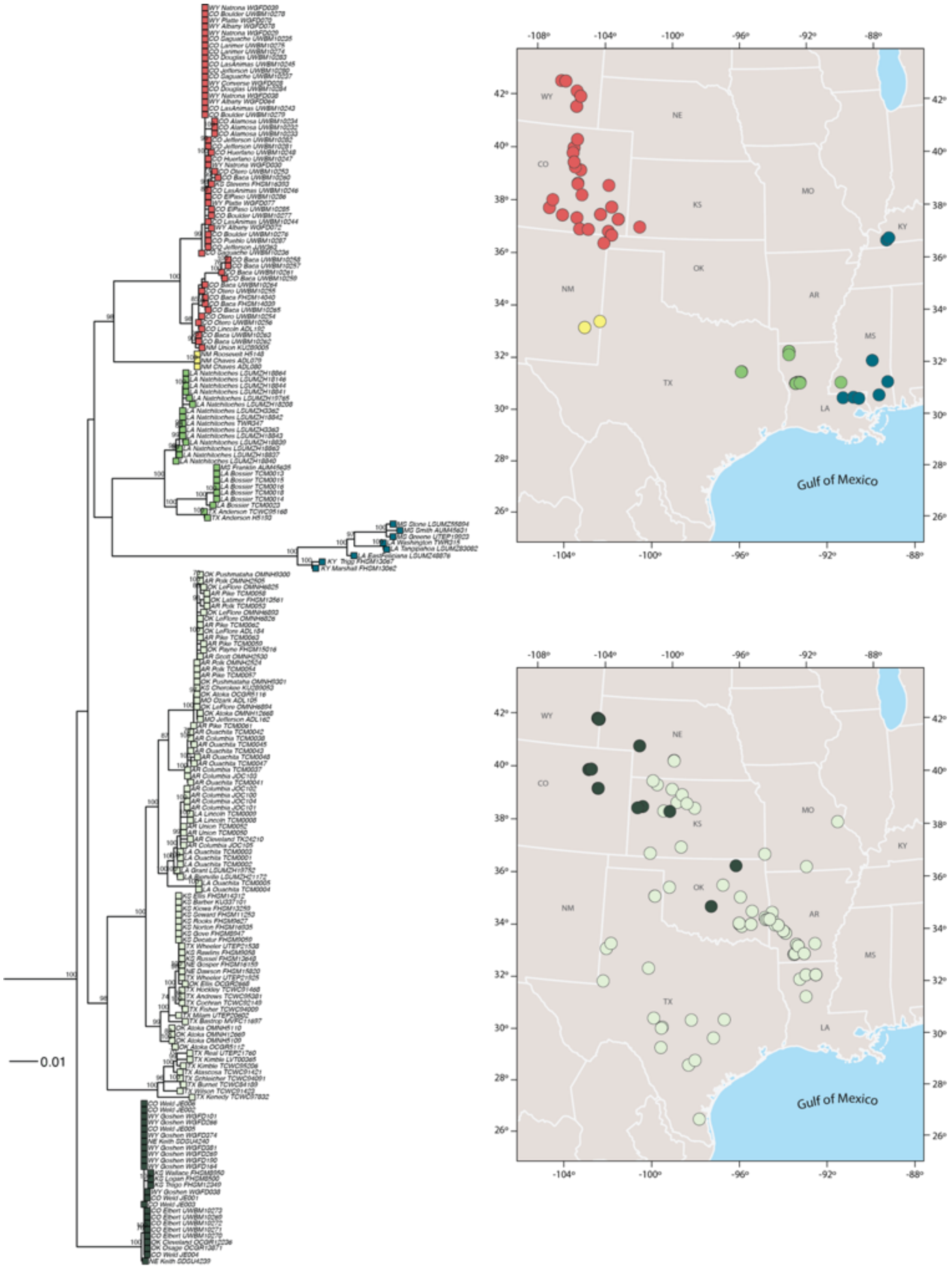


FIGURE S4. Phylogeographic patterns within *Sceloporus consobrinus* inferred using mtDNA. The phylogeny was estimated with IQTREE. Tips with the same colors reflect species delimitation results from the locMin method implemented in the R package Spider.

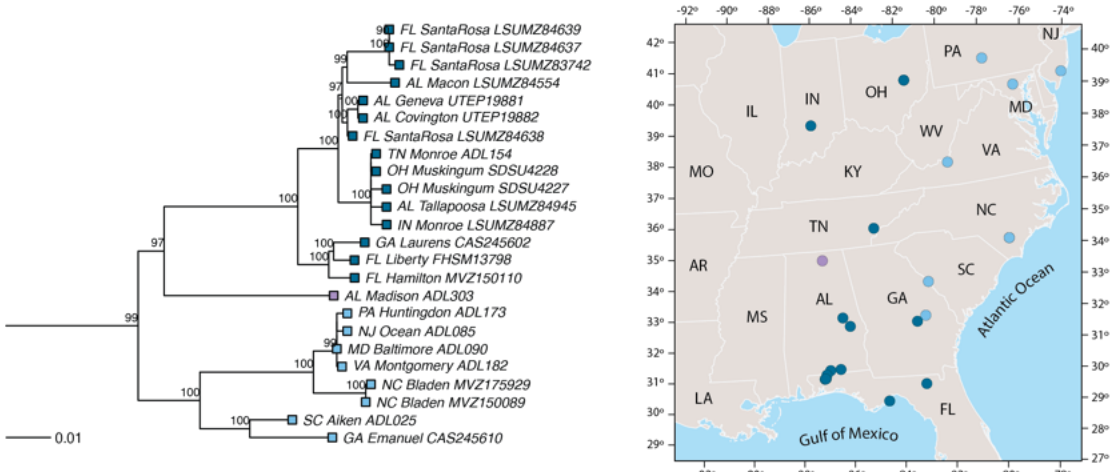


FIGURE S5. Phylogeographic patterns within *Sceloporus undulatus* inferred using mtDNA. The phylogeny was estimated with IQTREE. Tips with the same colors reflect species delimitation results from the locMin method implemented in the R package Spider.

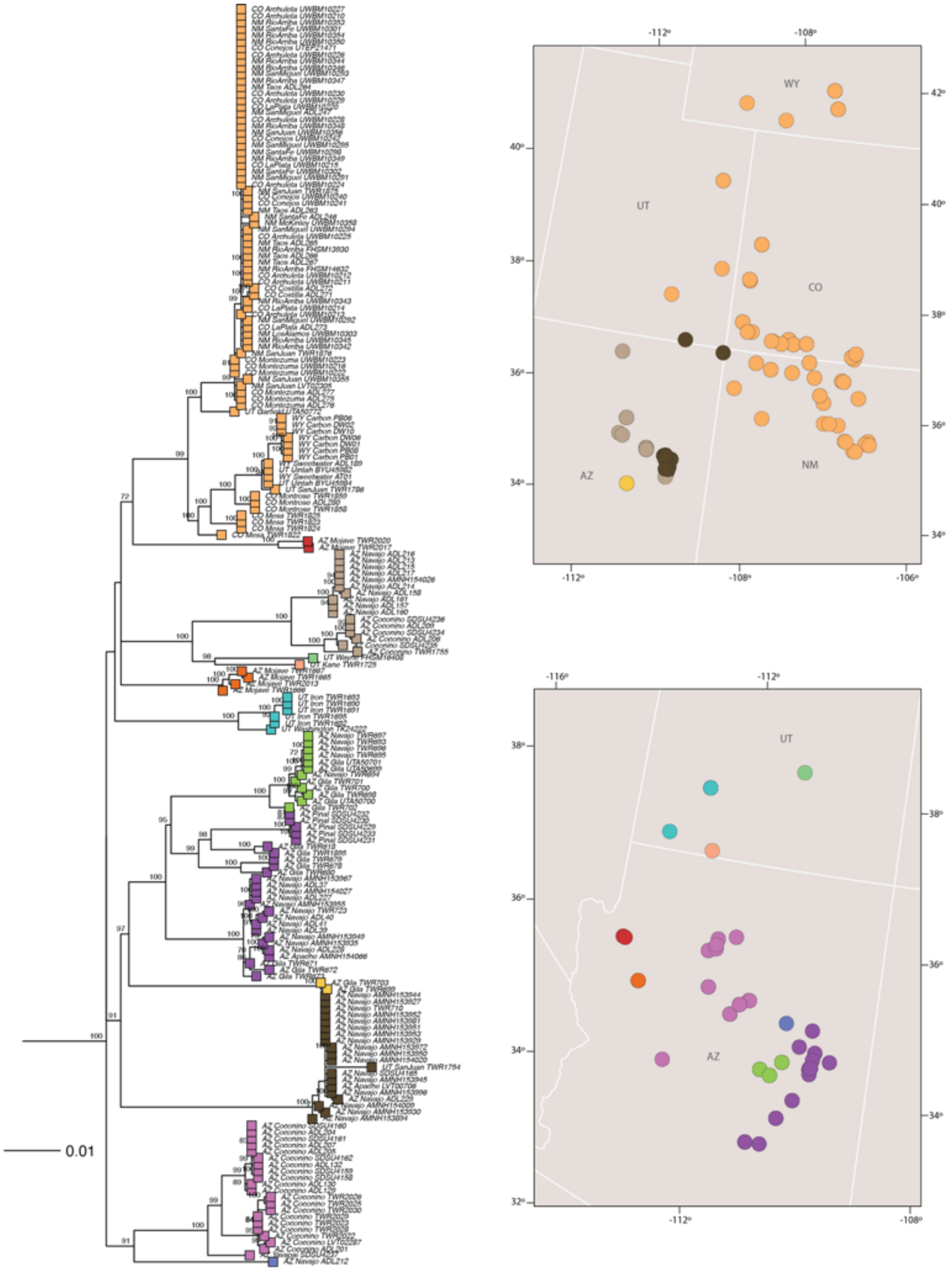


FIGURE S6. Phylogeographic patterns within *Sceloporus tristichus* inferred using mtDNA. The phylogeny was estimated with IQTREE. Tips with the same colors reflect species delimitation results from the locMin method implemented in the R package Spider.

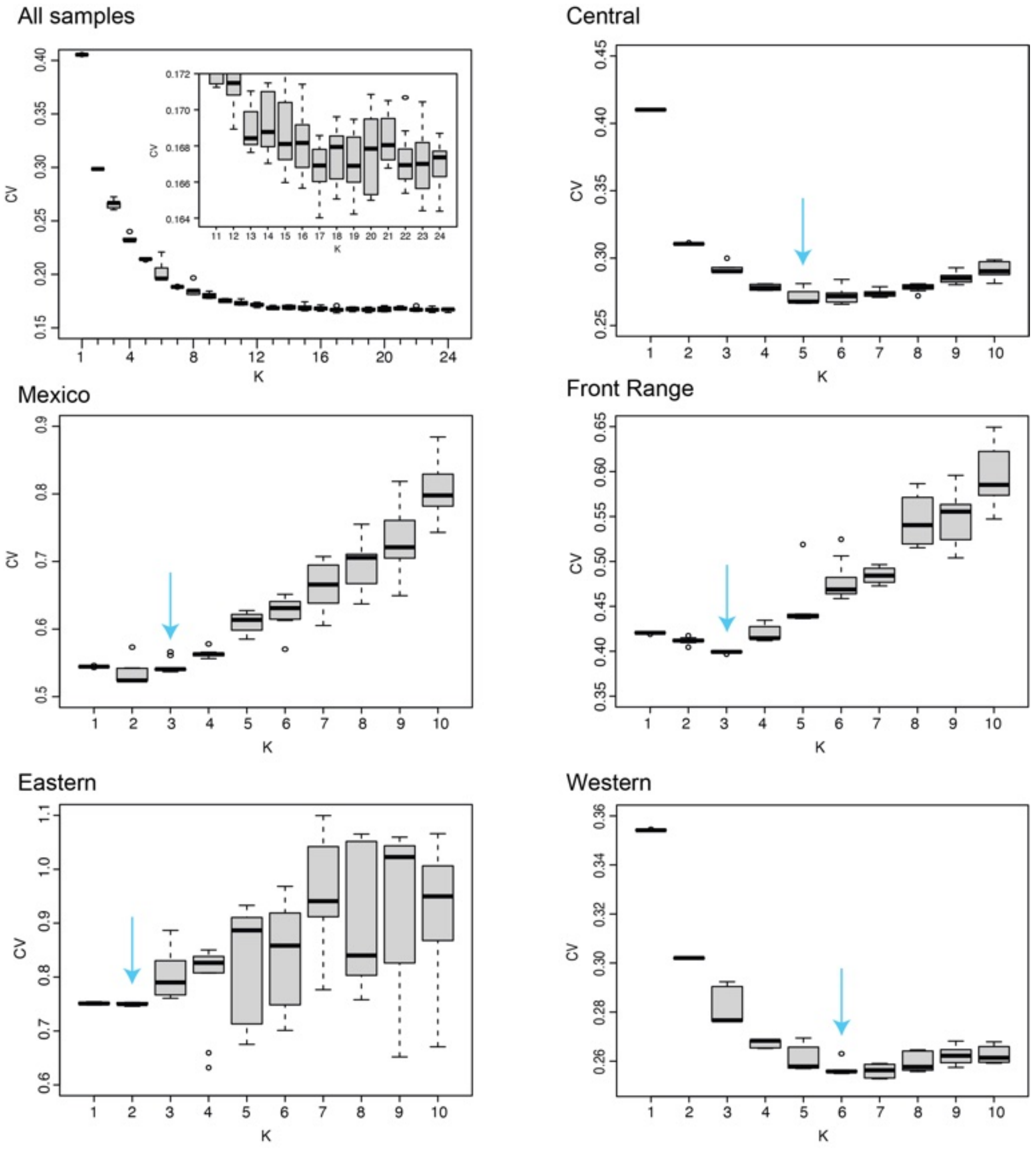


FIGURE S7. Admixture analysis results comparing the cross-validation (CV) error scores for each major clade and the analysis with all species combined (= all samples). The model with the combination of the lowest average CV error score and least variation across the 10 replicate runs was selected as the optimal model (indicated with an arrow).

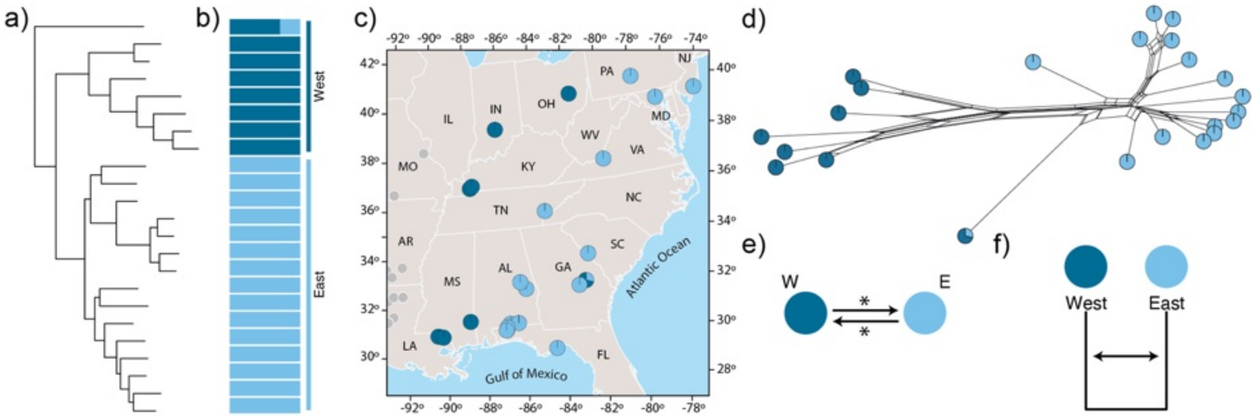


FIGURE S8. Phylogeography of the Eastern clade using nuclear data. a) Phylogeny estimated with ML using the concatenated data. b) Bar plot showing population structure ($K$ = 2). c) Geographic distributions of populations with pie charts showing admixture proportions. Gray dots represent samples belonging to other clades. d) Phylogenetic network with tips labeled by admixture proportions. e) Gene flow model based on the geographic connectivity of parapatric populations. Asterisk (*) = significant migration rate using Bayes factors. f) Species tree model used for migration estimation showing significant migration results with arrows.

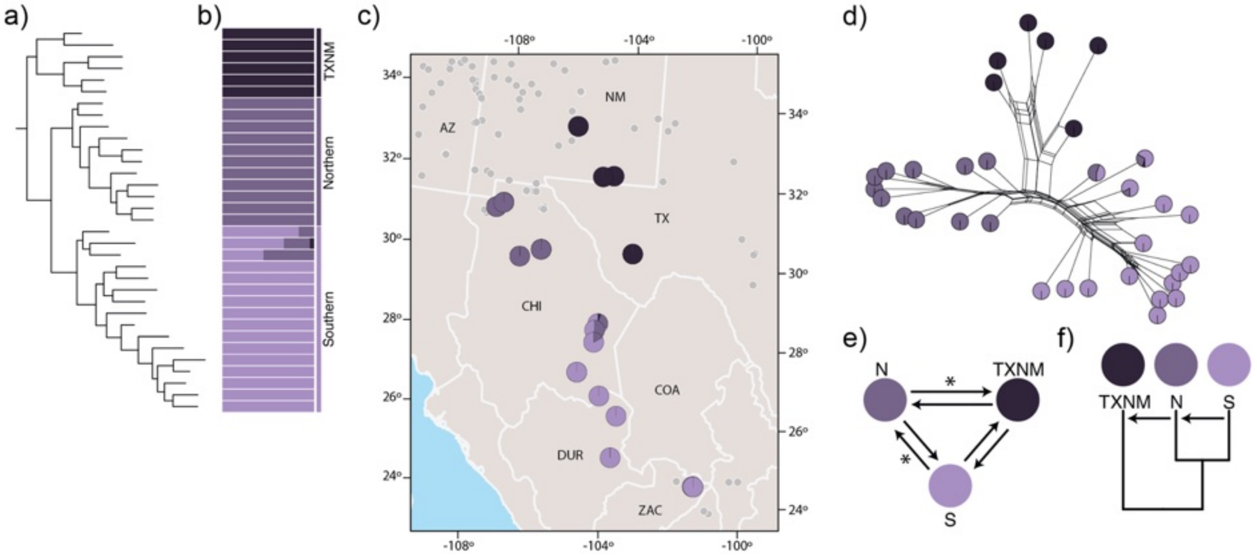


FIGURE S9. Phylogeography of the Mexico clade using nuclear data. a) Phylogeny estimated with ML using the concatenated data. b) Bar plot showing population structure (*K* = 3). c) Geographic distributions of populations with pie charts showing admixture proportions. Gray dots represent samples belonging to other clades. d) Phylogenetic network with tips labeled by admixture proportions. e) Gene flow model based on the geographic connectivity of parapatric populations. Asterisk (*) = significant migration rate using Bayes factors. f) Species tree model used for migration estimation showing significant migration results with arrows.

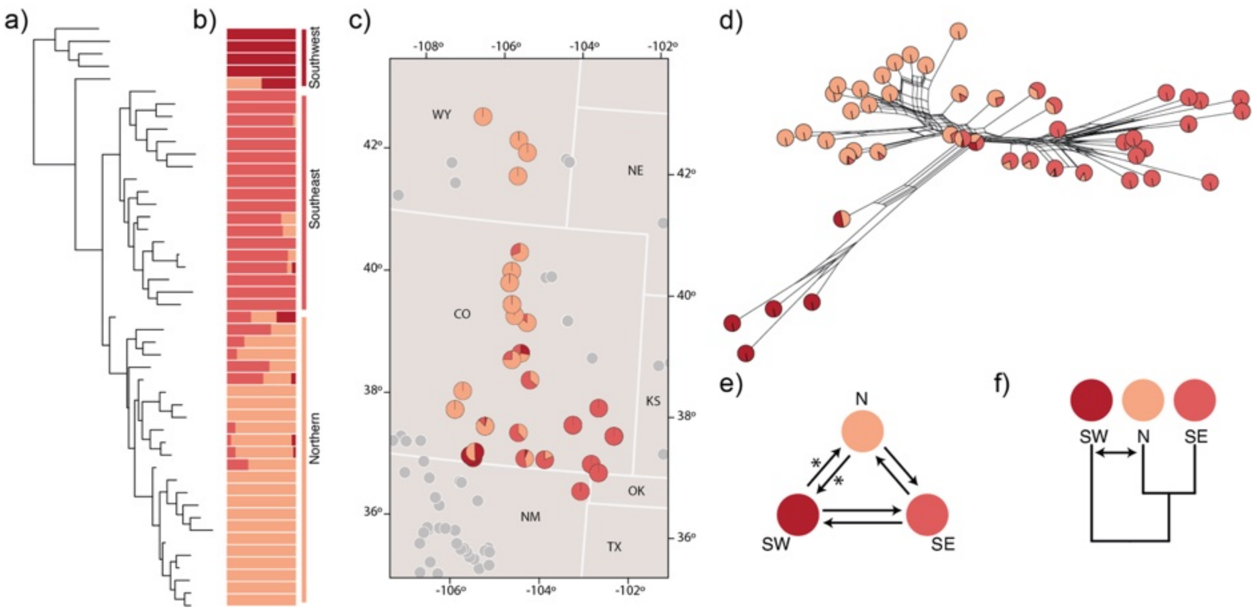


FIGURE S10. Phylogeography of the Front Range clade using nuclear data. a) Phylogeny estimated with ML using the concatenated data. b) Bar plot showing population structure (*K* = 3). c) Geographic distributions of populations with pie charts showing admixture proportions. Gray dots represent samples belonging to other clades. d) Phylogenetic network with tips labeled by admixture proportions. e) Gene flow model based on the geographic connectivity of parapatric populations. Asterisk (*) = significant migration rate using Bayes factors. f) Species tree model used for migration estimation showing significant migration results with arrows.

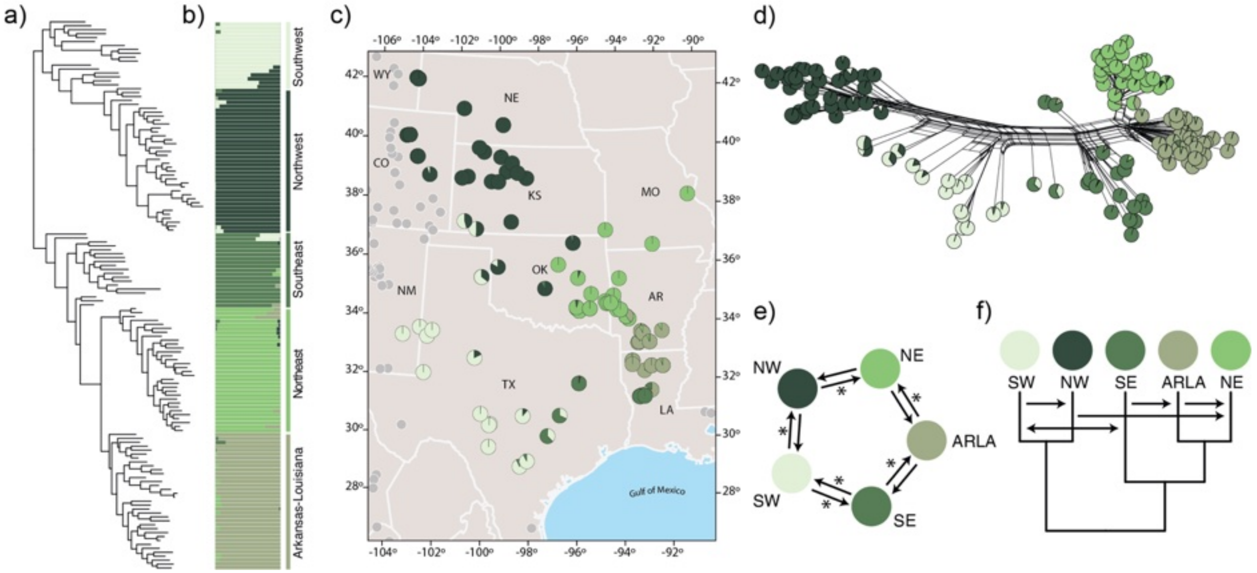


FIGURE S11. Phylogeography of the Central clade using nuclear data. a) Phylogeny estimated with ML using the concatenated data. b) Bar plot showing population structure (*K* = 5). c) Geographic distributions of populations with pie charts showing admixture proportions. Gray dots represent samples belonging to other clades. d) Phylogenetic network with tips labeled by admixture proportions. e) Gene flow model based on the geographic connectivity of parapatric populations. Asterisk (*) = significant migration rate using Bayes factors. f) Species tree model used for migration estimation showing significant migration results with arrows.

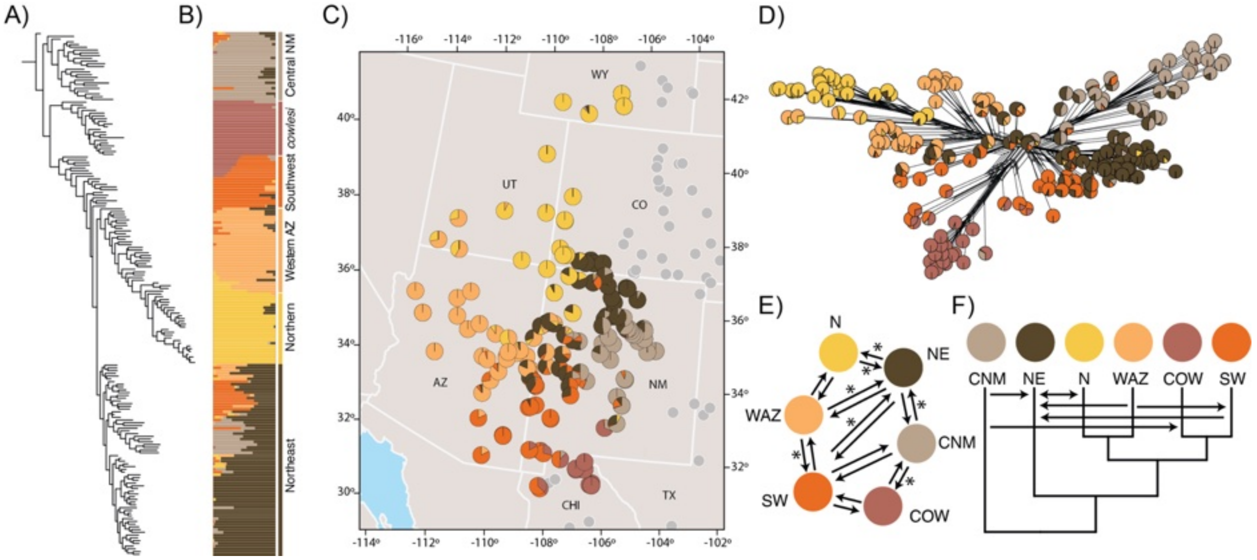


FIGURE S12. Phylogeography of the Western clade using nuclear data. a) Phylogeny estimated with ML using the concatenated data. b) Bar plot showing population structure ($K$ = 6). c) Geographic distributions of populations with pie charts showing admixture proportions. Gray dots represent samples belonging to other clades. d) Phylogenetic network with tips labeled by admixture proportions. e) Gene flow model based on the geographic connectivity of parapatric populations. Asterisk (*) = significant migration rate using Bayes factors. f) Species tree model used for migration estimation showing significant migration results with arrows.

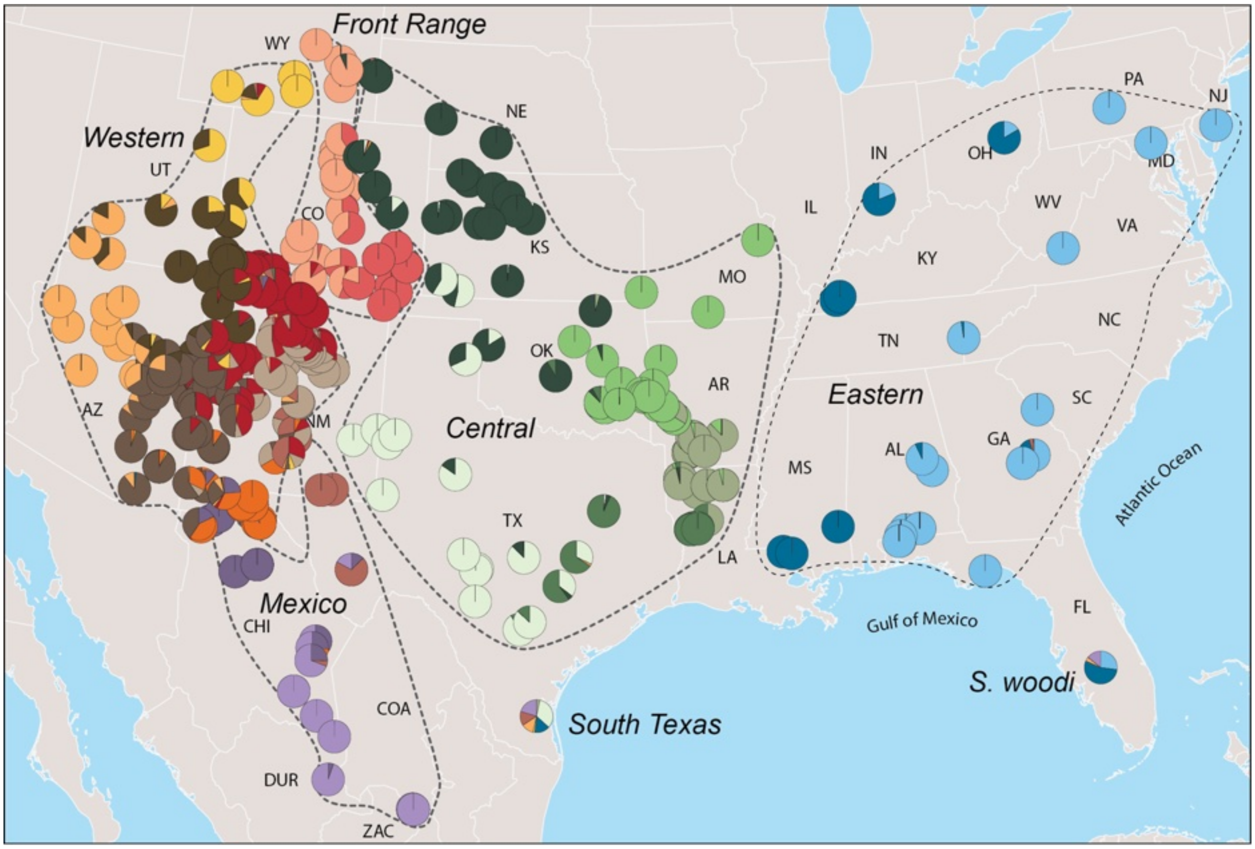


FIGURE S13. Admixture results ($K$ = 19) for the analysis of all samples in the *undulatus* complex. Pie charts show admixture proportions. Dotted lines circumscribe major groups. The geographic distribution of samples in the western portion of the range are shown in detail in Fig. S14. Instead of representing distinct populations, the two divergent lineages represented by single samples (e.g., *S. woodi* and South Texas) are admixed with other populations.

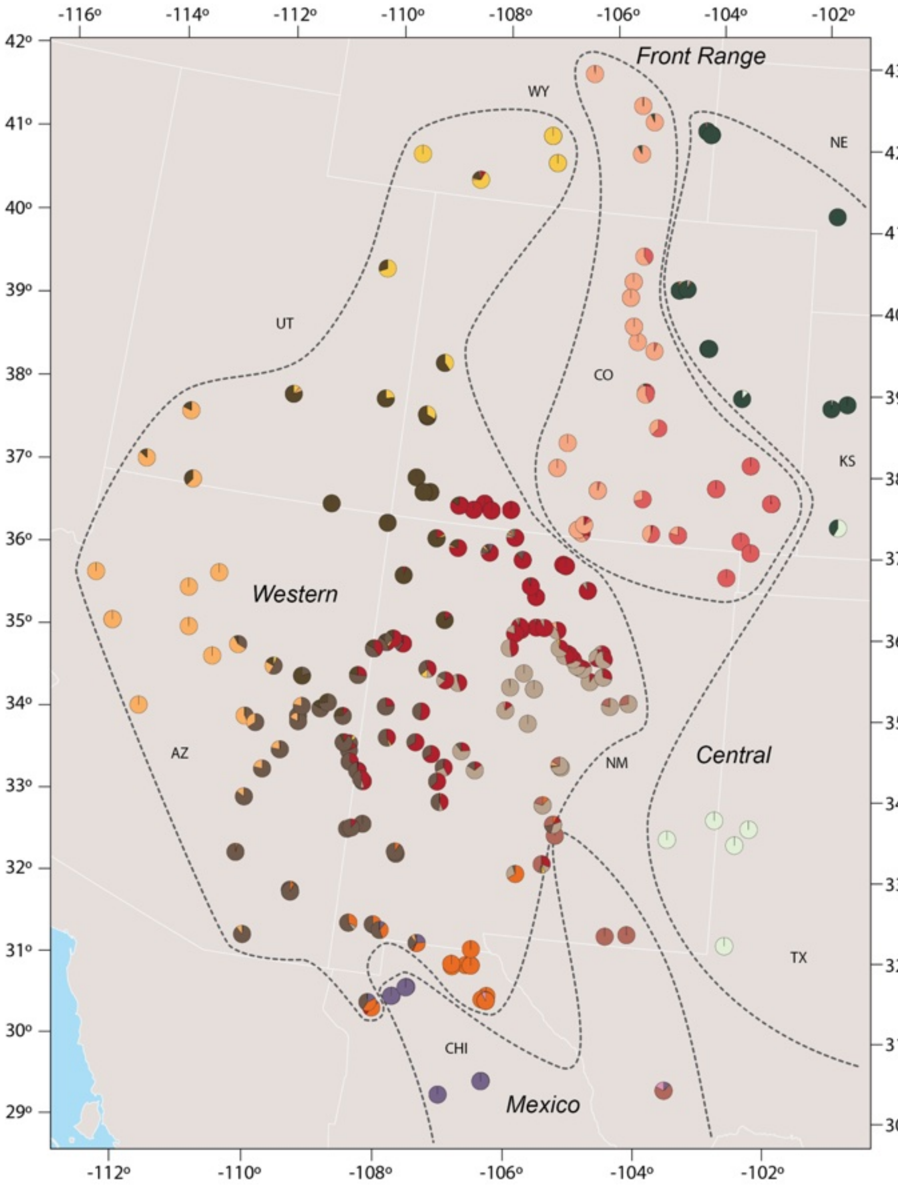


FIGURE S14. Detailed view of admixture results (*K* = 19) for the western distribution of the *undulatus* complex. Pie charts show admixture proportions. Dotted lines circumscribe major groups.

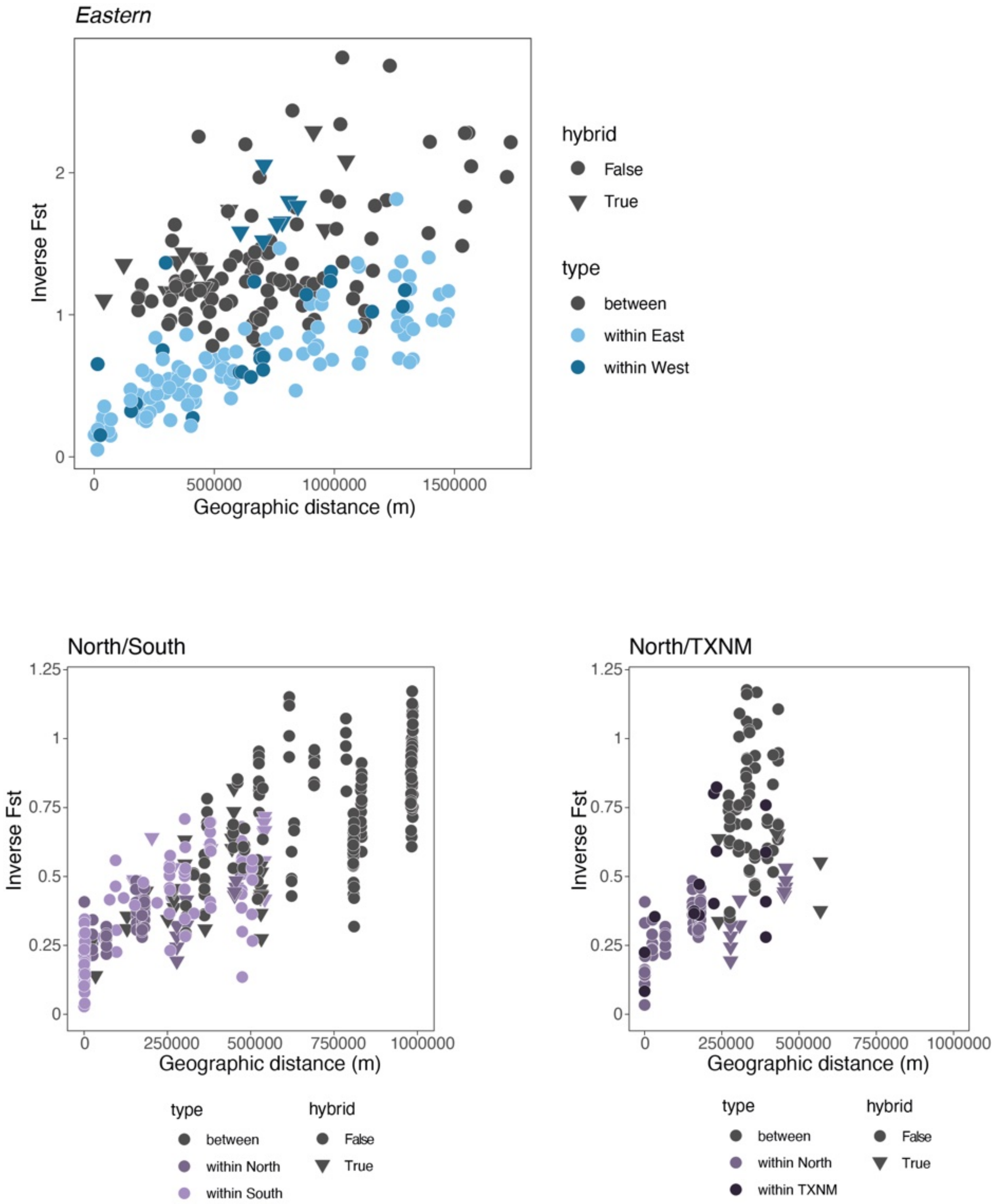


FIGURE S15. Isolation-by-distance in the *Sceloporus undulatus* complex. Eastern (top) and Mexico (bottom). Hybrid individuals are those with > 20% probability of ancestry in two or more groups.

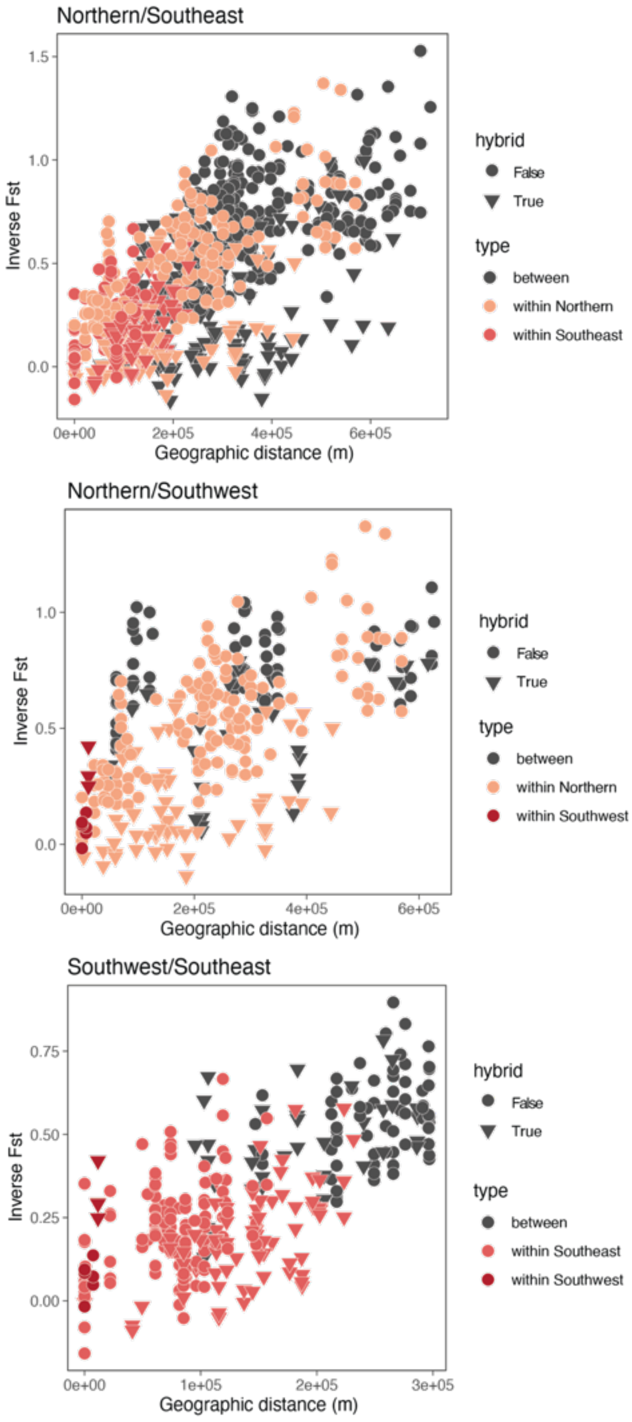


FIGURE S16. Isolation-by-distance in the Front Range clade. Hybrid individuals are those with > 20% probability of ancestry in two or more groups.

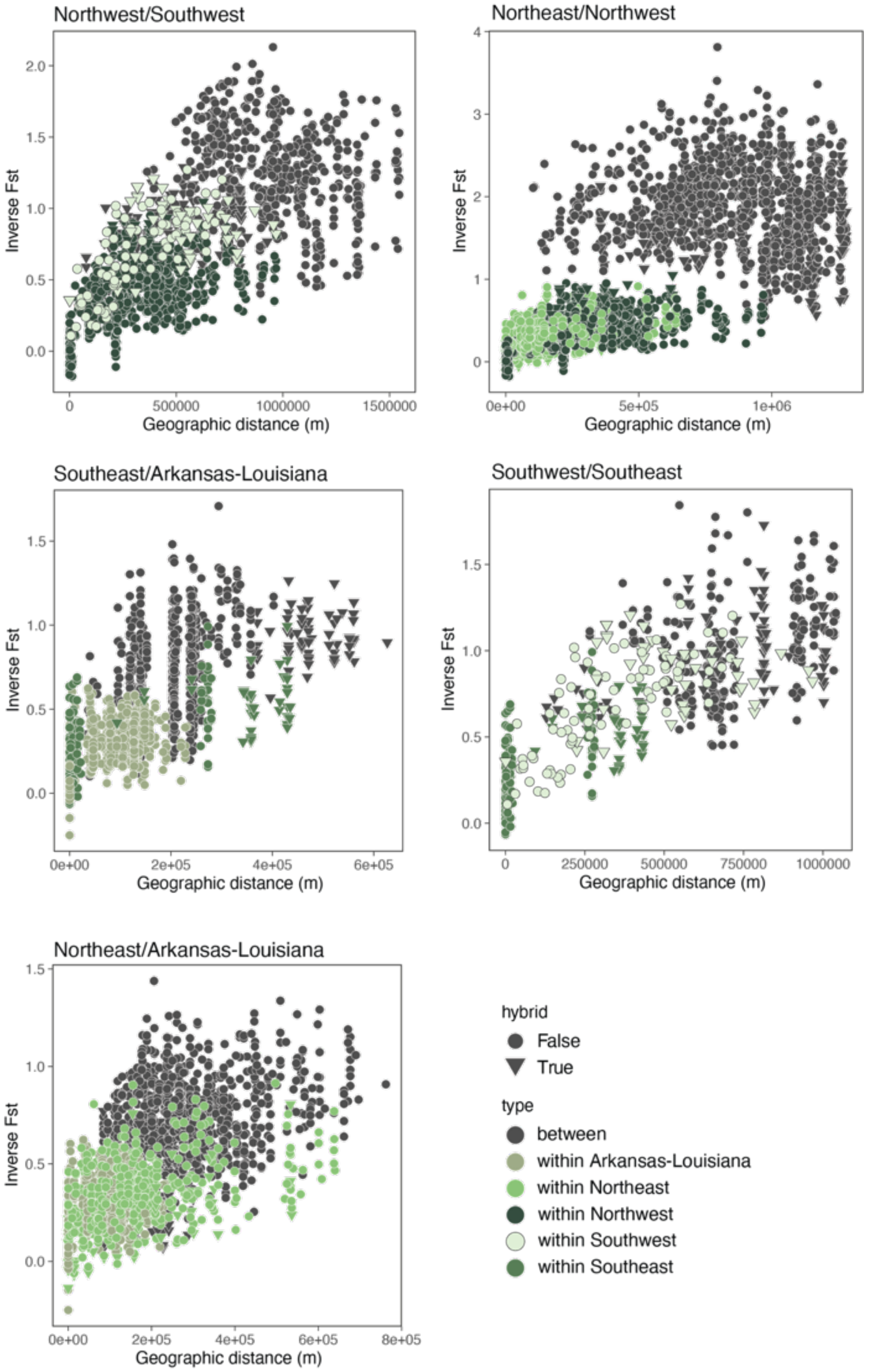


FIGURE S17. Isolation-by-distance in the Central clade. Hybrid individuals are those with > 20% probability of ancestry in two or more groups.

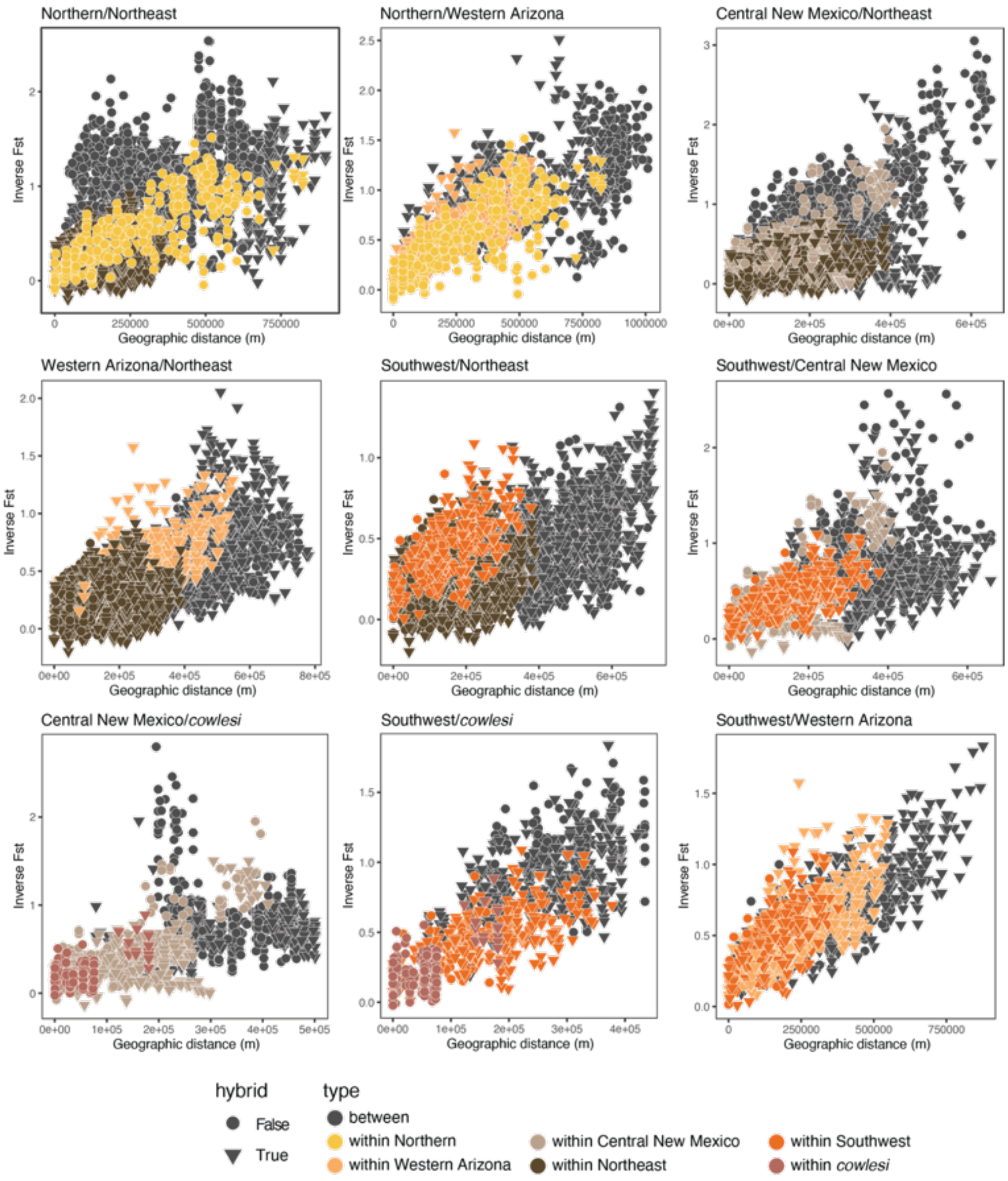


FIGURE S18. Isolation-by-distance in the Western clade. Hybrid individuals are those with > 20% probability of ancestry in two or more groups.

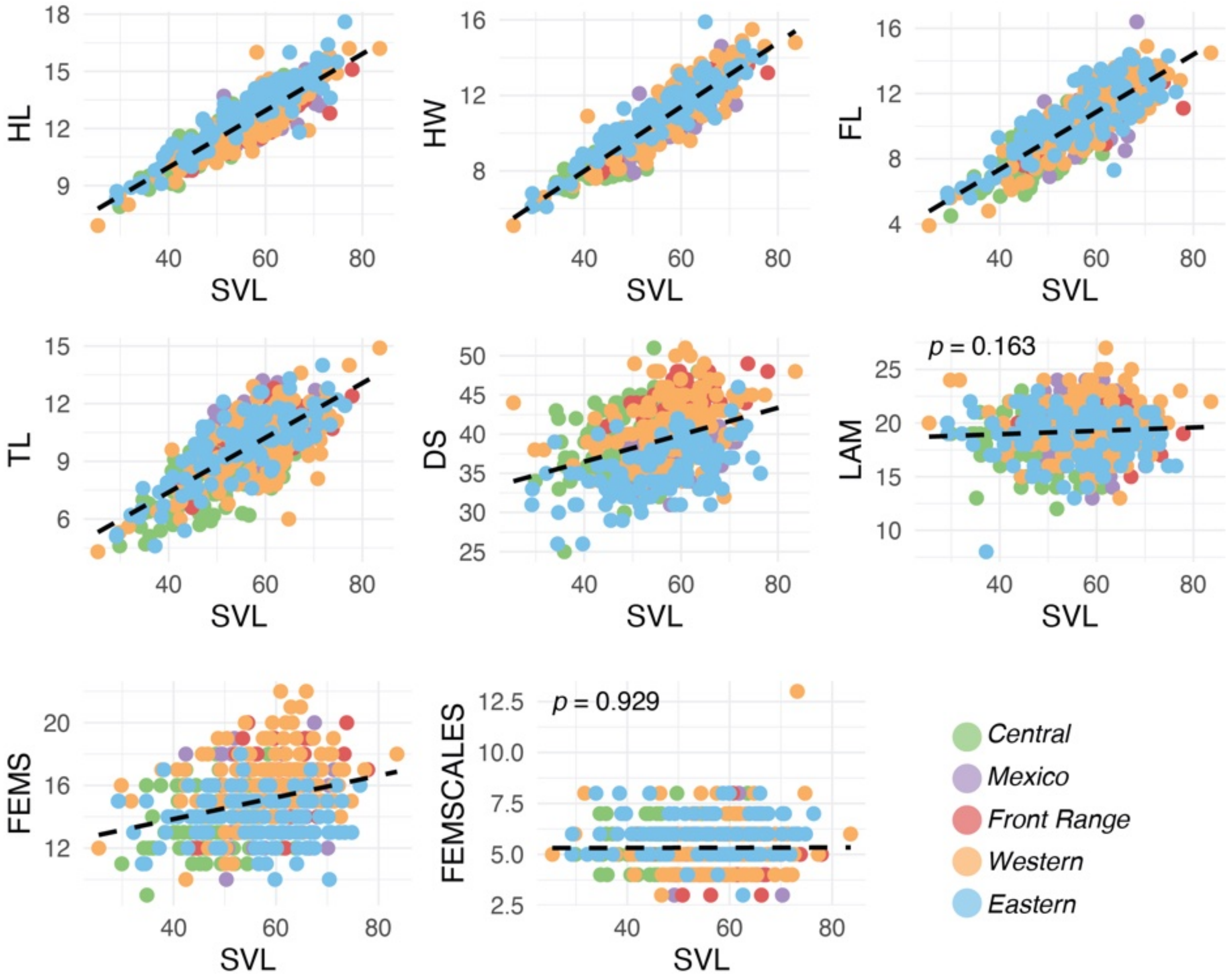


FIGURE S19. Scatterplots showing relationships between traits and body size (SVL). Linear models showed significant correlations for all traits except LAM and FEMSCALES.

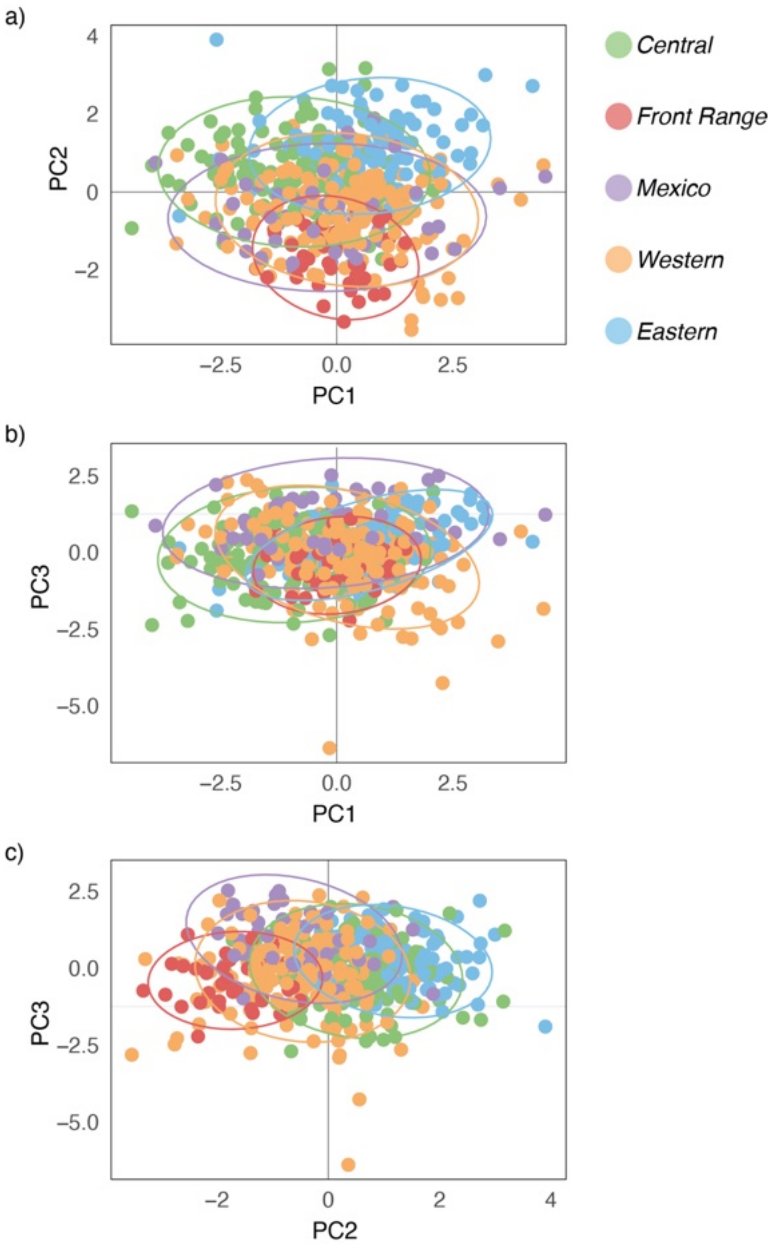


FIGURE S20. PCA of the morphological data. A) PC1 vs. PC2. B) PC1 vs. PC3. C) PC2 vs. PC3. Genetic groups are color-coded. The first three components account for 60% of the variation.

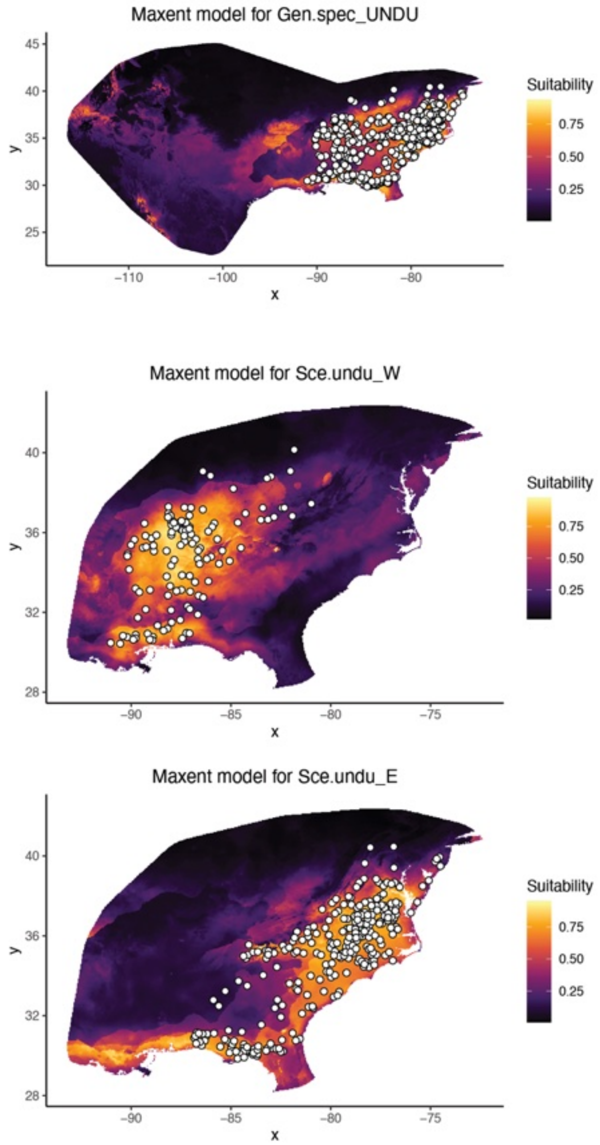


FIGURE S21. Ecological niche models for the Eastern clade and two populations within the clade. Dots indicate localities used to generate the models. Warmer colors indicate a higher probability for species presence.

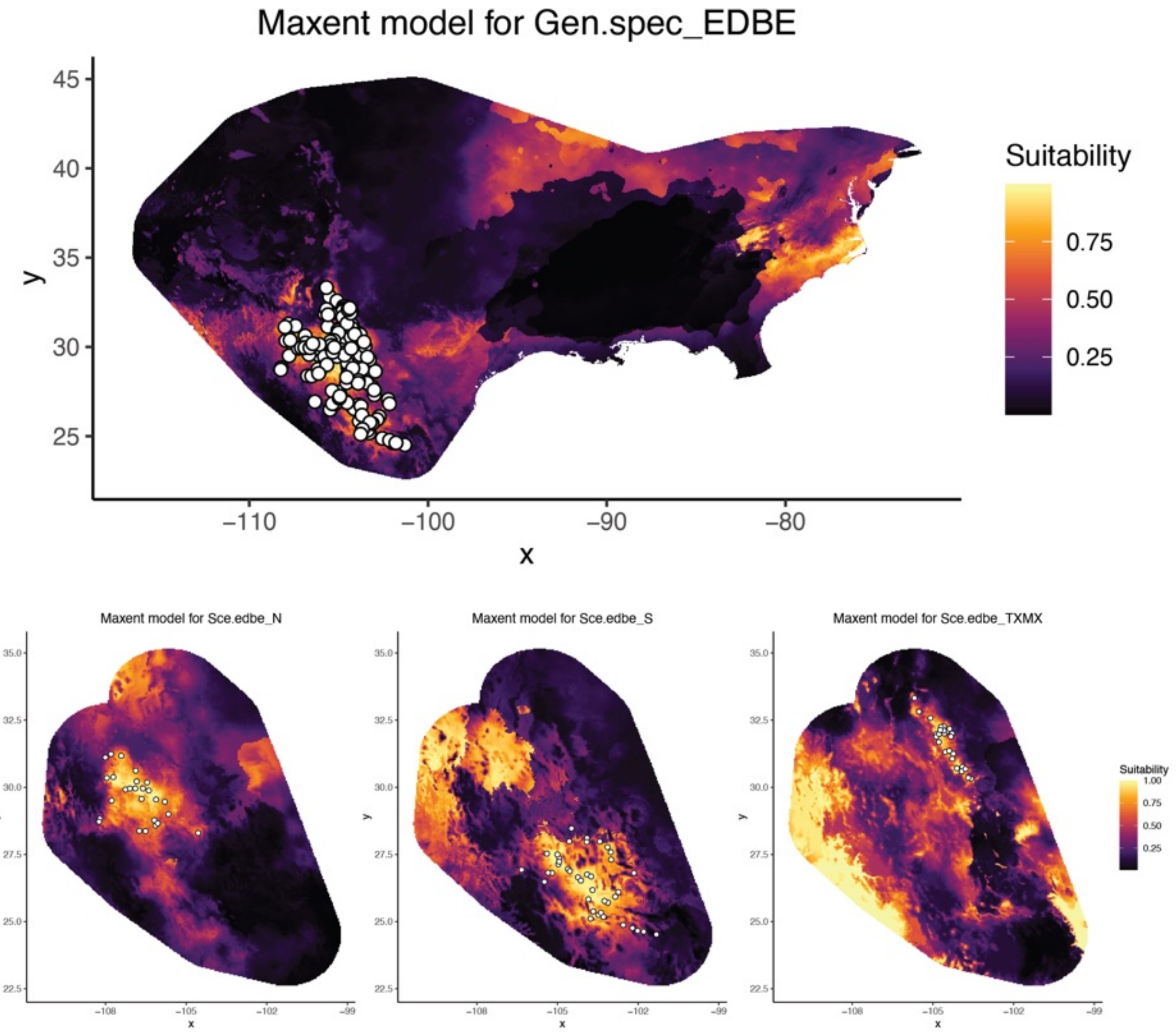


FIGURE S22. Ecological niche models for the Mexico clade and three populations within the clade. Dots indicate localities used to generate the models. Warmer colors indicate a higher probability for species presence.

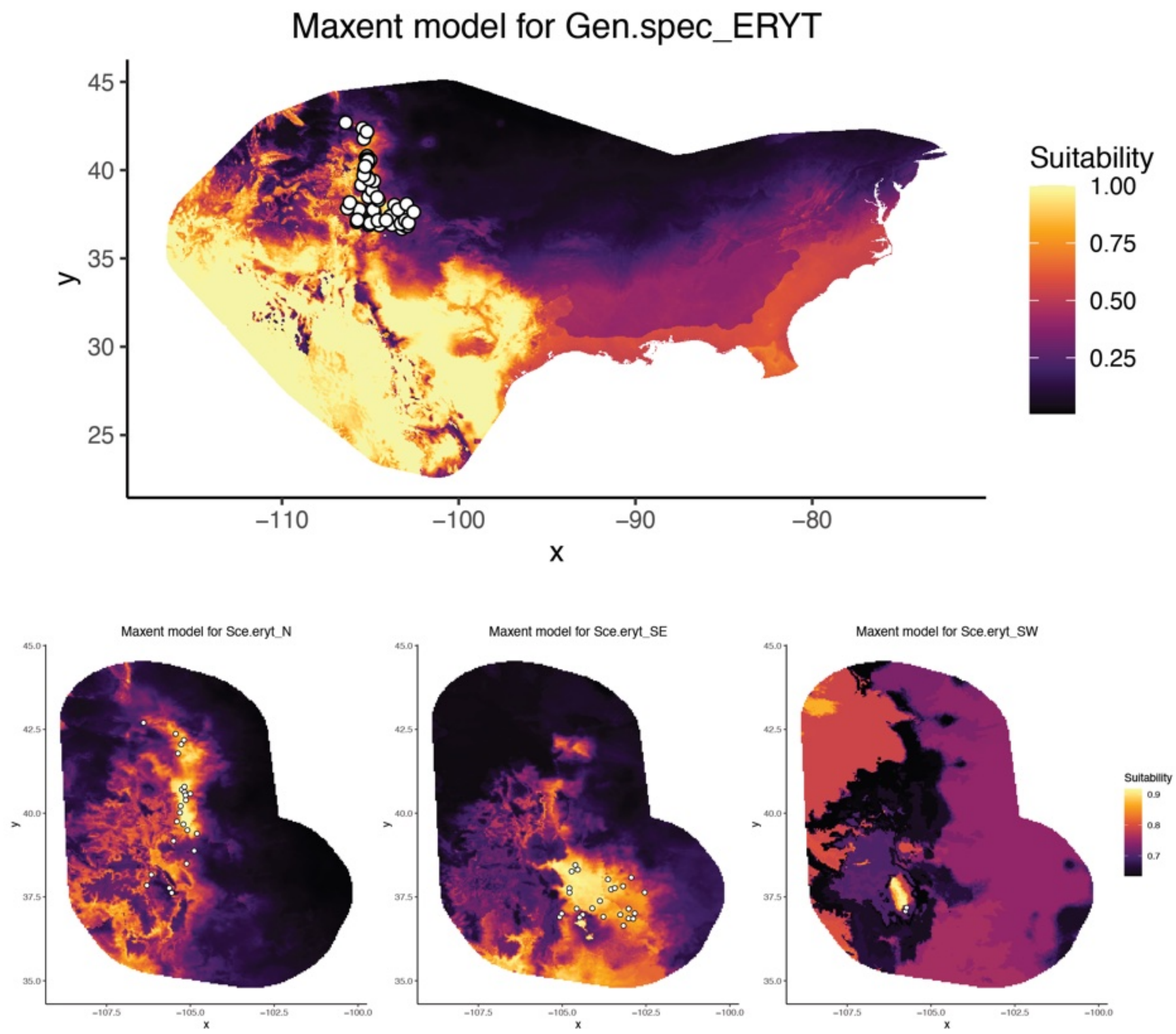


FIGURE S23. Ecological niche models for the Front Range clade and three populations within the clade. Dots indicate localities used to generate the models. Warmer colors indicate a higher probability for species presence.

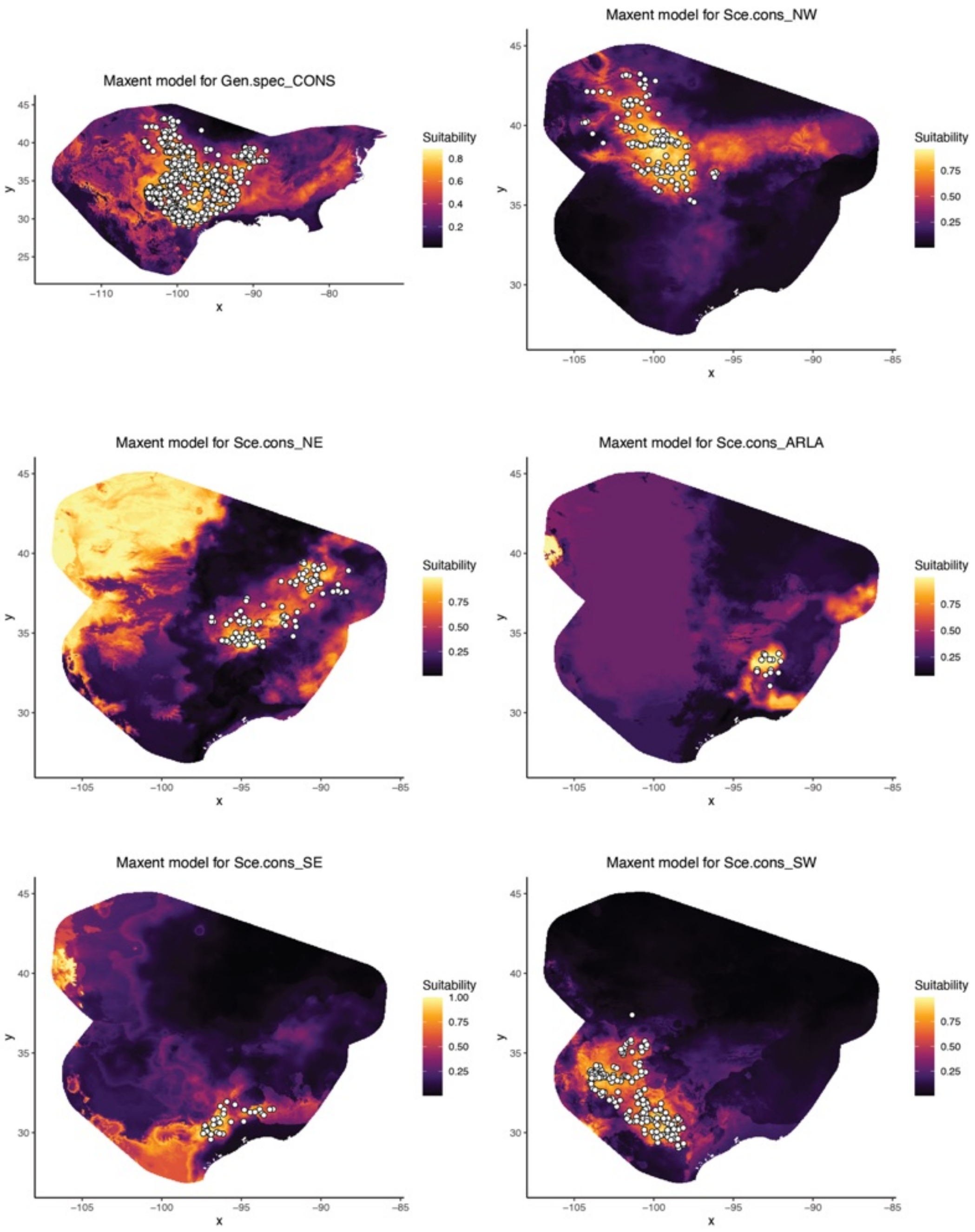


FIGURE S24. Ecological niche models for the Central clade and five populations within the clade. Dots indicate localities used to generate the models. Warmer colors indicate a higher probability for species presence.

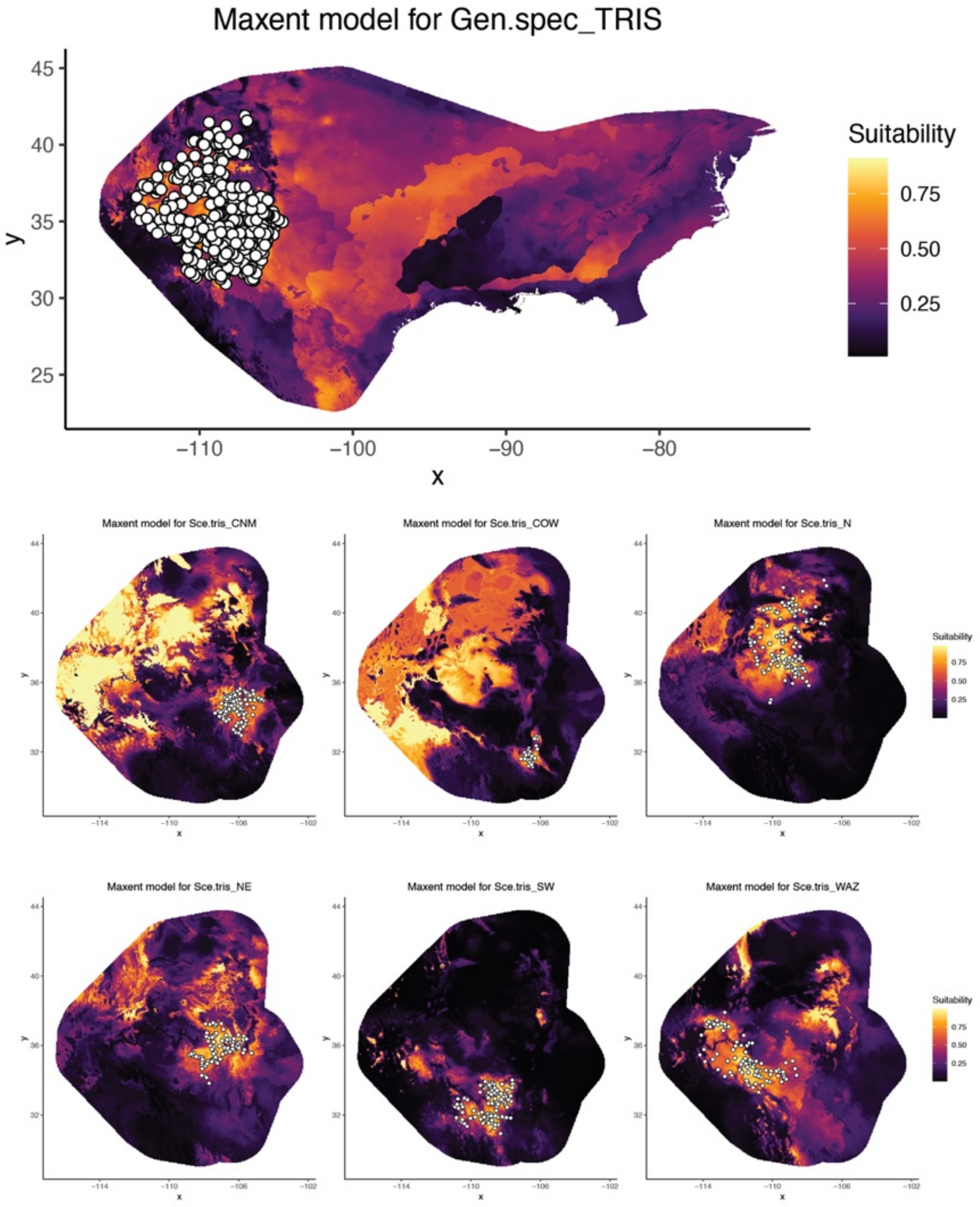


FIGURE S25. Ecological niche models for the Western clade and six populations within the clade. Dots indicate localities used to generate the models. Warmer colors indicate a higher probability for species presence.

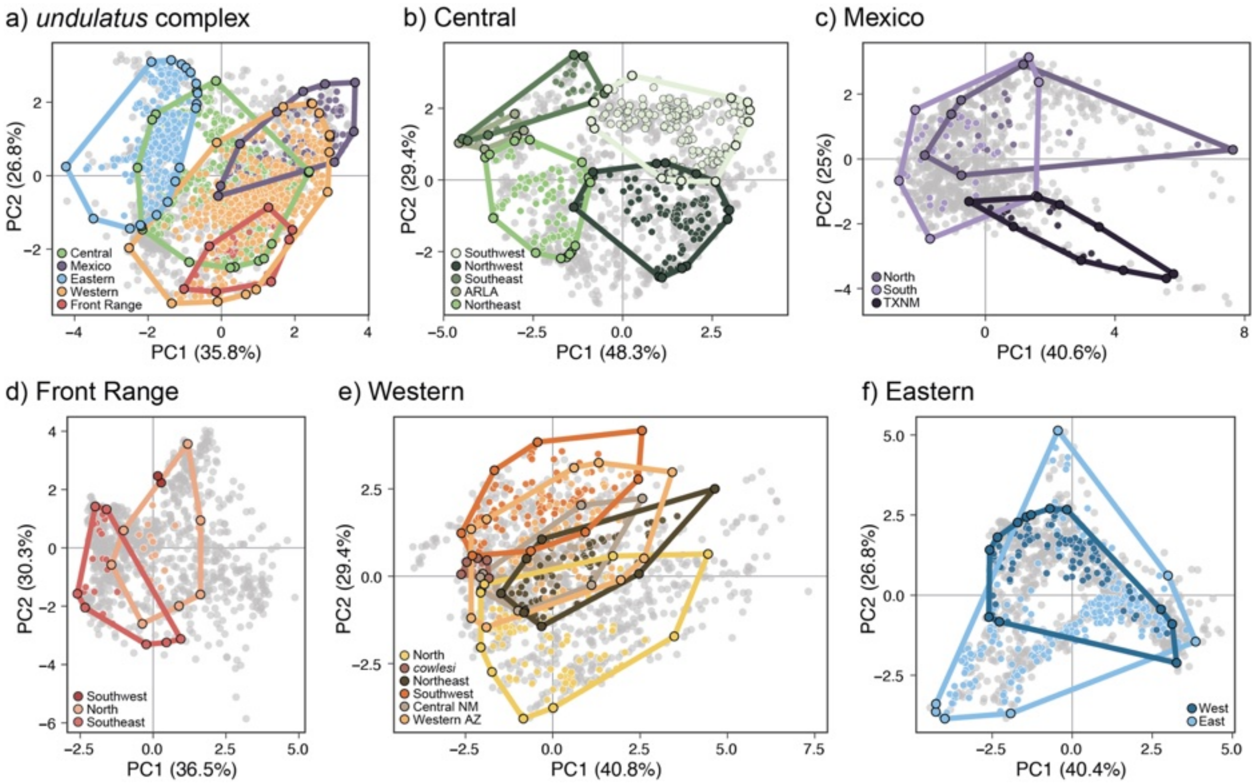


FIGURE 26. Principal component analysis of bioclimatic variables and environmental space for the *Sceloporus undulatus* complex. Colored points represent occurrence records assigned to populations and colored lines show the minimum convex polygon (MCP) encompassing all occurrences. Grey dots show the background environment defined as the MCP of all known occurrences plus a 200 km buffer, sampled with 1,000 random points. a) Broad-scale analysis of the five major widespread clades in the *undulatus* complex, b) Central clade, c) Mexico clade, d) Front Range, e) Western clade, f) Eastern clade.

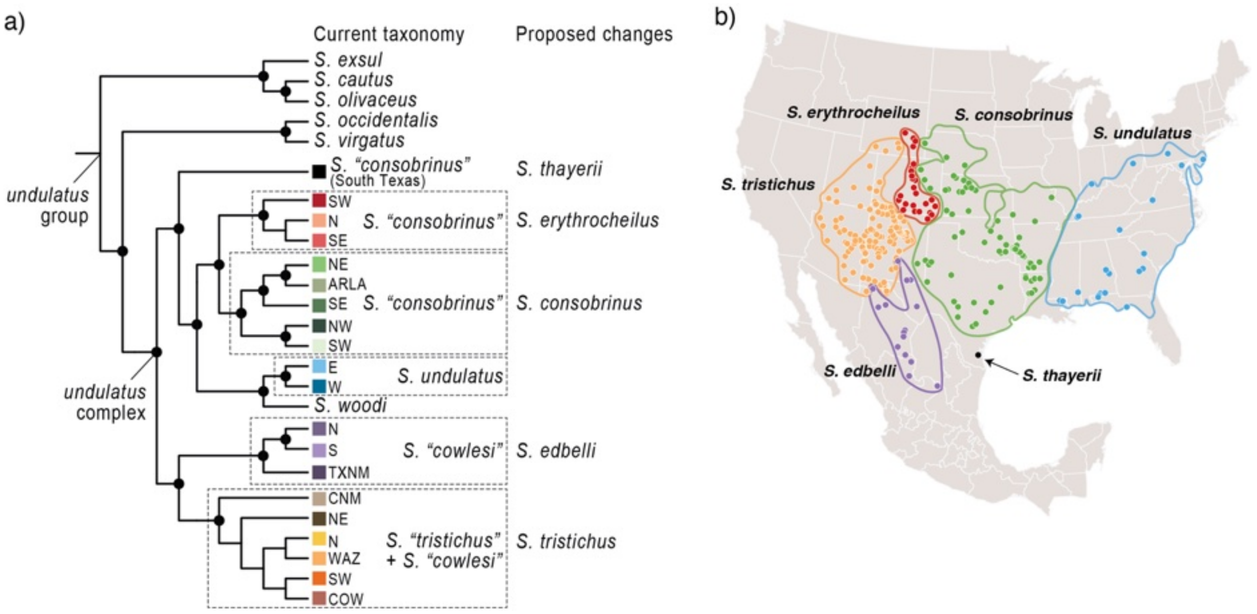


FIGURE S27. Phylogenetic relationships in the *Sceloporus undulatus* group based on the concatenated ddRADseq data analyzed in IQTREE. The current taxonomy is based on the mtDNA. Proposed taxonomic changes to the *undulatus* complex are illustrated next to the phylogeny and shown on the map.

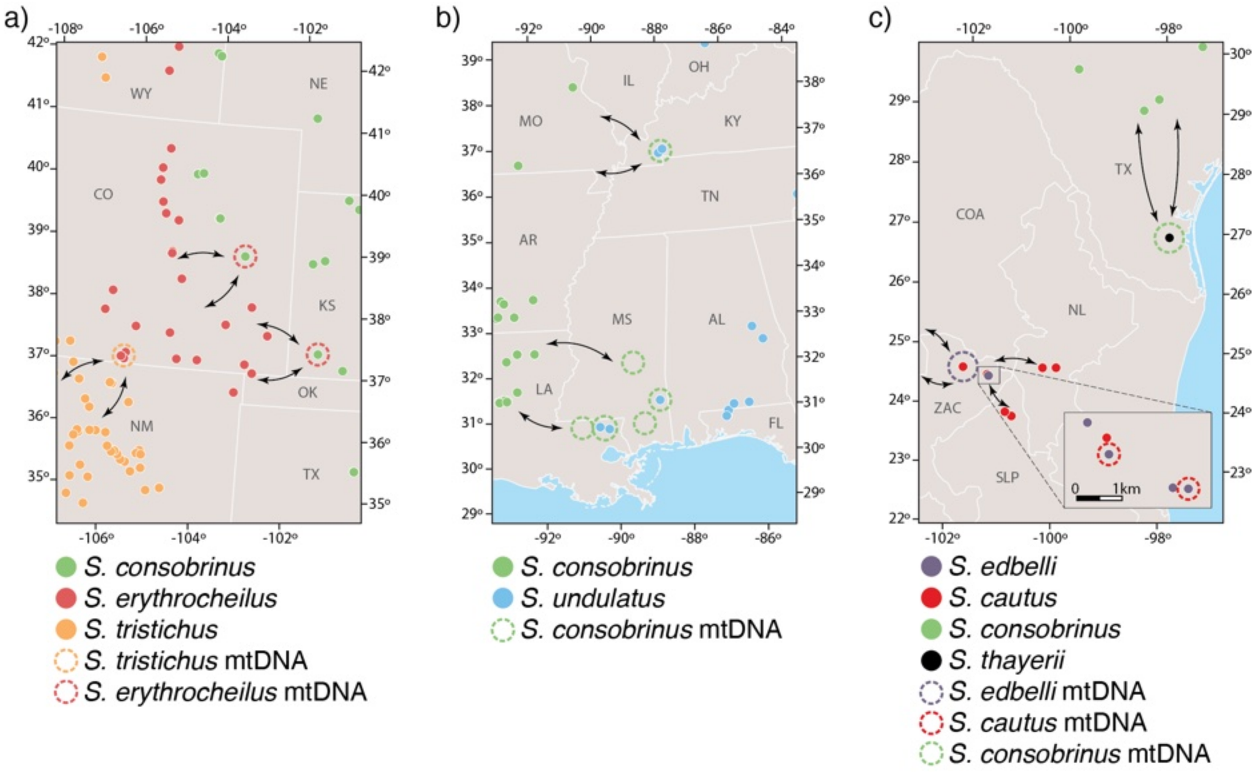


FIGURE S28. Mitochondrial introgression traces to the contact zones between five adjacent species pairs. Maps show the geographic distributions of species based on nuclear DNA. Introgression is illustrated with open circles, and arrows connect the interacting species pairs. a) mtDNA belonging to *S. tristichus* is found in southwestern populations of *S. erythrocheilus,* and *S. erythrocheilus* mtDNA occurs in *S. consobrinus.* b) mtDNA from *S. consobrinus* is found in western populations of *S. undulatus.* Introgression is inferred for several of the western *S. undulatus* populations based on their geographic proximity to other *S. undulatus*. c) *Sceloporus thayerii* has a mtDNA haplotype belonging to *S. consobrinus.* Examples of mtDNA introgression between *S. cautus* and *S. ebelli* occur in both directions.
